# HER2; p53 Co-mutated Cancers Show Increased Histone Acetylation and are Sensitive to Neratinib plus Trastuzumab Deruxtecan

**DOI:** 10.1101/2025.07.06.663368

**Authors:** Xiaoqing Cheng, Jacob Hsia, Jesus Iraheta, Joanna Gongora, Maureen Highkin, Xiaohua Jin, Zhanfang Guo, Julie L. Prior, John R. Edwards, Shunqiang Li, Ian S. Hagemann, Cynthia X. Ma, Zongtao Lin, Benjamin A. Garcia, Ron Bose

## Abstract

In metastatic breast cancer, *HER2*-activating mutations often co-occur with *TP53* mutations, a combination linked to poor response to neratinib and worse prognosis. To model this clinical challenge, we bred *HER2* ^V777L^ transgenic mice with two *TP53* mutant alleles: *TP53* ^R172H^ (the murine homolog of human *TP53* R175H) and *TP53*^fl/fl^, which mimics p53 truncations common in human tumors. *TP53* mutations accelerated tumor development and reduced survival in *HER2*-mutant mice. These co-mutant tumors were resistant to neratinib but remained sensitive to exatecan, the topoisomerase I (TOP1) inhibitor payload in trastuzumab deruxtecan (T-DXd). Mechanistically, *TP53* mutant tumors exhibited upregulation of histone acetylation, hypertranscription of DNA repair factors, increased chromatin accessibility, and rendered cells more susceptible to TOP1 inhibitors via G2/M arrest and apoptosis. This vulnerability is dependent on transcriptional activity of *TP53* mutations, highlighting a novel strategy to treat *HER2;TP53* co-mutant breast cancers using TOP1-targeted therapies.

**Statement of Significance:** *TP53* mutations sensitize HER2-mutant cancers to TOP1 inhibitors via chromatin accessibility and hyper-transcription, supporting combination therapy with neratinib and T-DXd in *TP53*/HER2 co-mutant breast cancers.

## Introduction

Human epidermal growth factor receptor 2 (HER2) is a member of the EGFR receptor tyrosine kinase family. It plays a crucial role in cell proliferation and survival and is a major drug target in the treatment of breast cancer and other carcinomas. ^1, 2^ *HER2* gene amplification or overexpression was found in about 15-20% of breast cancer, and in other cancers, such as gastric, ovarian, and endometrial cancers. HER2 activating mutations are specific genetic changes in the *HER2* gene that cause ligand-independent activation of the HER2 signaling pathway without gene amplification or protein overexpression. These mutations produce constant (constitutive) activation of HER2’s tyrosine kinase domain, triggering downstream signaling, contributing to tumor formation and metastasis ^1, 3–5^. The MutHER and SUMMIT clinical trials showed that neratinib could be used to treat patients with HER2-activating mutations^6, 7^. These trials found that patients whose breast cancers were co-mutated in *HER2* and *TP53* had lower objective response rates and shorter progression-free survival than *HER2*-mutated cancer patients without *TP53* mutations ^8, 9^. *TP53* co-mutation occurs in a significant fraction (30–60%) of HER2-mutant tumors, especially in breast cancer^3^, non-small cell lung cancer (NSCLC)^10^, and bladder cancer.^11^

*TP53* acts as a genome instability guardian, and the mutation frequency in breast cancer varies. 30-40% of HER2 mutated breast cancers have P53 mutations, and 70-80% of triple-negative (ER-, PR-, HER2-) breast cancers contain P53 mutations.^12^ Mutant *TP53* is considered an oncogene through “separation-of-function” at the transcriptional level, promoting chromosomal instability^13^ and unveiling therapeutic opportunities.^14^ 50% of human cancers carry *TP53* mutations and 80% of *TP53* mutations are located in the DNA binding domain^15^. In some cases, hotspot mutations in the p53 DNA binding domain are common and give the mutant p53 gain-of-function properties, while truncating mutations are less common, leading to a completely nonfunctional p53 protein, either through frameshift mutations or nonsense mutations, resulting in loss of function activity. While some mouse models expressing p53^R172H^ and p53^R270H^ mutations (mutations at codons 175 and 273 in humans, respectively) in lung^16^, colon^17^, pancreas^18, 19^, and intestinal^20^ tissues have been shown to develop adenocarcinomas, p53^R172H^ and p53^R270H^ mice also developed novel patterns of tumors compared to non-mutated p53(p53KO) mice, including sarcomas^21, 22^, lymphomas^23, 24^ and more frequent endothelial tumors^20, 25^, indicating a different mechanism of tumorigenesis between disrupted *TP53* and its gain of function mutations ^26^. The knowledge gap in the mechanism and contributions of *TP53* mutations to drug response activity challenges therapeutic targets of *TP53* and results in an undruggable target in this field.

Trastuzumab deruxtecan (T-DXd) is an antibody-drug conjugate (ADC) that delivers a cytotoxic payload derived from exatecan, a highly potent topoisomerase I (TOP1) inhibitor. Exatecan is a synthetic analog of camptothecin, a natural product that inhibits topoisomerase I, an enzyme critical for DNA replication^27^. TOP1 inhibitors are anticancer drugs, and are used to treat a range of solid tumors and hematologic malignancies by trapping the TOP1-DNA cleavage complex, the trapped DNA topoisomerases (TOPs) disrupt replication and transcription, leading to DNA damage and apoptosis.^28,29^ Trastuzumab deruxtecan is currently FDA-approved for the treatment of breast and gastric cancer, HER2-mutated non-small cell lung cancer and any HER2 overexpressing (HER2 IHC 3+) solid tumors ^30^.

To better understand the drug response activity role of p53 mutations in HER2-mutated cancer patients in the SUMMIT trial,^9^ and to clarify the different roles of mutations between PIK3CA and p53,^31^ this paper generated new co-mutated mouse models of breast cancer: HER2^V777L^; p53*^fl/fl^* (H53fl/fl) and HER2^V777L^; p53*^R172H^*(H53fl/172). We found that H53^fl/fl^ and H53^fl/172^ tumors with the *TP53* mutation accelerated tumorigenesis and progression. H53 co-mutated tumors had increased histone acetylation and chromatin accessibility, thereby sensitizing the HER2 and p53 co-mutated tumor cells to T-DXd. Combining neratinib and T-DXd in HER2; p53 co-mutated breast cancer was a very effective, pre-clinical treatment for these cancers. The neratinib and T-DXd combination is currently being tested in a phase I, multi-institutional clinical trial (NCI ETCTN 10495, ClinicalTrials.gov accession number NCT05372614).

## Results

### Mutated *TP53* accelerates tumorigenesis and shortens survival time

We bred our HER2 mutant transgenic mouse with two different alleles of *TP53*, p53fl/fl and p53R172H, in order to model what occurs in human breast cancer. About one-third of *TP53* mutations seen in human breast cancers are truncating mutations, which are represented in our experiments by the *TP53*fl/fl allele. The other two-thirds of *TP53* mutations seen in human breast cancers are missense mutations, typically in the DNA binding domain, and they are represented in our experiments by mice with the *TP53*R172H allele (murine *TP53*R172H is the human homolog of *TP53*R175H mutation). Further, *TP53* missense mutations can also co-occur with loss of the wild-type allele, termed Loss of Heterozygosity (LOH). To model these three possibilities in HER2 mutant breast cancer, we bred the genotypes of mice indicated in Fig. 1A. The generation of these different genotypes with different *TP53* alleles was necessary because we found that these genotypes affected the response to trastuzumab deruxtecan (DS-8201a; T-DXd), which utilizes a DNA-damaging chemotherapy moiety. We have already developed a transgenic mouse that conditionally expresses the human HER2 V777L cDNA (abbreviated as “H”), which was reported previously^31, 32^. We bred H mice with Rosa26-LSL-p53fl/fl and Rosa26-LSL-p53R172H mutant transgenic mice to get the heterozygous (abbreviated as H53wt/fl, H53wt/172) and homozygous p53 (abbreviated as H53fl/fl, H53fl/172) co-mutant with H. Orthotopic and intraductal injection of adenovirus expressing Cre (Ad-Cre) into 10-week-old female mice caused Cre recombinase-mediated removal of the lox-STOP-lox cassette, ^31^ generating mammary tumors in these mice (Fig. 1B). H mice slowly developed areas of ductal *in situ* carcinomas and invasive adenocarcinomas over a 7–10-month period after Ad-Cre injection (Fig. 1B). H53fl/fl mice developed tumors with sarcomatoid carcinoma, and H53fl/172 mice developed adenocarcinoma, confirmed by a pathologist. Multiple immunostains were performed on the carcinomas from H53 mutant mice to characterize the mammary biomarker status of H53 tumors. The H53 tumors are ER-positive, HER2-positive, and smooth muscle actin (SMA) negative (Fig. 1C).

**Fig. 1:**
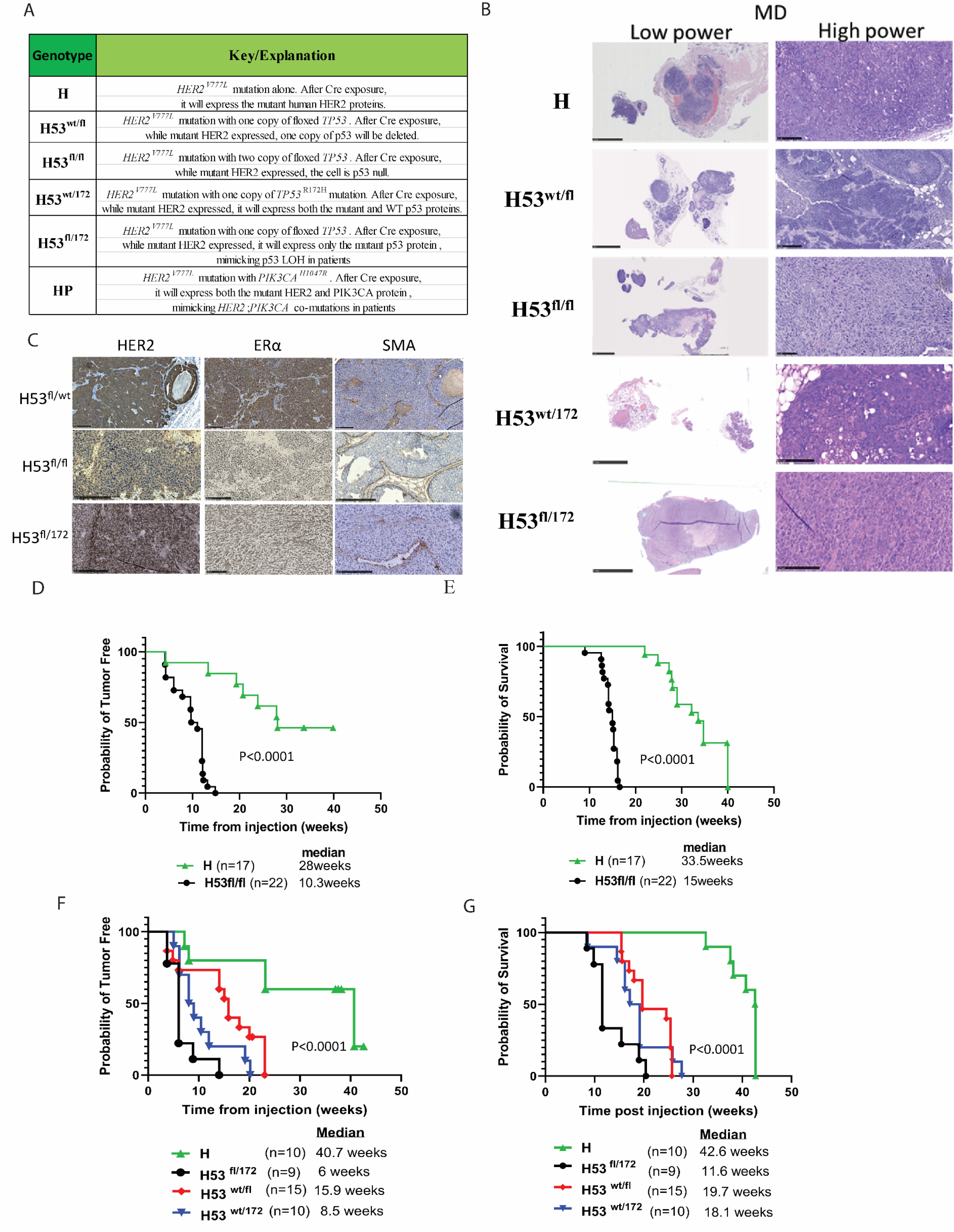
Characterization of H53 tumors in mice with HER2V777L and P53fl/fl or P53R172H. (A) Key to the genotype of the murine breast cancer models. (B) Representative hematoxylin and eosin (H&E) staining of image of tumor slides from H, H53wt/fl, H53wt/172, H53fl/fl, H53fl/172 mice. Scale bars of low-power images are 5 mm. Scale bars of high-power images are 500 µm. (C) Immunohistochemistry (IHC) staining of HER2, ER*α*, SMA of mammary gland tumor tissue sections from H53wt/fl, H53fl/fl, and H53fl/172 mice, respectively. Scale bars of low-power images are 5 mm. Scale bars of high-power images are 100 µm. (D) Kaplan-Meier analysis shows overall survival (right) and tumor-free (left) of H, and H53 null mice, respectively. (E) Kaplan-Meier analysis shows overall survival (right) and tumor-free (left) of H, H53wt/fl, H53wt/172, and H53fl/172 mice, respectively.

We first compared the rate of tumor formation in the H53fl/fl mice to that of our prior mice that only carried the HER2 mutation. Our previous work showed HP mice formed tumors the fastest (median of 2.1 weeks after injection with Adeno-Cre) ^31^, followed by a median for tumor formation of 10 weeks in H53fl/fl mice and a median tumor formation of 28 weeks in HER2 mutant-only mice (Fig. 1D). Similarly, median overall survival was three weeks for HP mice (previously reported), 15 weeks for H53fl/fl mice, and 33 weeks for H mice (Fig. 1E). We then examined other *TP53* genotypes in our transgenic mouse model and found that H53fl/172 was the fastest to form tumors (Fig. 1F) and kill the mice (Fig. 1G; median of 6 weeks and 11.6 weeks, respectively), followed by H53wt/172 (median tumor-free period 8.5 weeks and median survival of 18 weeks).

Together, these data indicate that H53 mutant tumors mimic and display human breast cancer features; p53 mutations accelerate tumorigenesis and shorten the survival of the H53 co-mutant mice.

### Organoids derived from H53 tumors show resistance to Neratinib and are more sensitive to TOP1 inhibitors *in vitro*

Organoids isolated from the H53^fl/fl^ and H53^fl/172^ showed more budding morphology, which is considered the feature of aggressive growth in organoid culture, than the cells derived from H, H53^wt/fl^, and H53^wt/172^ mice (Fig. 2A). Mammary biomarker status of H53 tumor organoid was validated by immunohistochemistry staining (Fig. 2B). To confirm that these specific deficiencies in the *TP53* and HER2 co-mutant cancer cells are associated with clinical/drug sensitivity responses, we tested the effect of the different *TP53* genotypes on neratinib sensitivity and found that H53fl/fl and H53fl/172 organoids were the most resistant to neratinib (Fig. 2C). H53wt/172 organoids also showed resistance to neratinib, though not as pronounced as H53fl/fl and H53fl/172. In contrast, the order of drug sensitivity to exatecan was very different and could varied experiment to experiment (Fig. 2D). We found that most H53fl/fl experiments were 10-30-fold more sensitive to exatecan. In contrast, H53wt/172 and H53fl/172 organoids were 5-7-fold more resistant to exatecan (Fig. 2D), in keeping with the general consensus that *TP53* mutations cause resistance to chemotherapy. Variability between independent biological samples of the same genotype suggests that secondary events occurring in tumor formation, perhaps due to genome instability caused by *TP53* alterations, could be contributing to the varying sensitivity to exatecan. Exatecan selectively inhibits TOP1 by trapping the TOP1-DNA reaction, the cleavage complex. The toxicity addresses the advantage of targeting macromolecules. The most plausible mechanism to explain the efficiency of targeting TOP1-DNA complexes in cancer cells is the deficiency of DNA repair and cell cycle checkpoints in cases where severe DNA damage results due to inhibited TOP1 causing polymerases to be stuck in both replication and transcription. Cell cycle arrest causes mitotic catastrophe upon TOP1 inhibition (Supplemental Fig. 1).

**Fig. 2:**
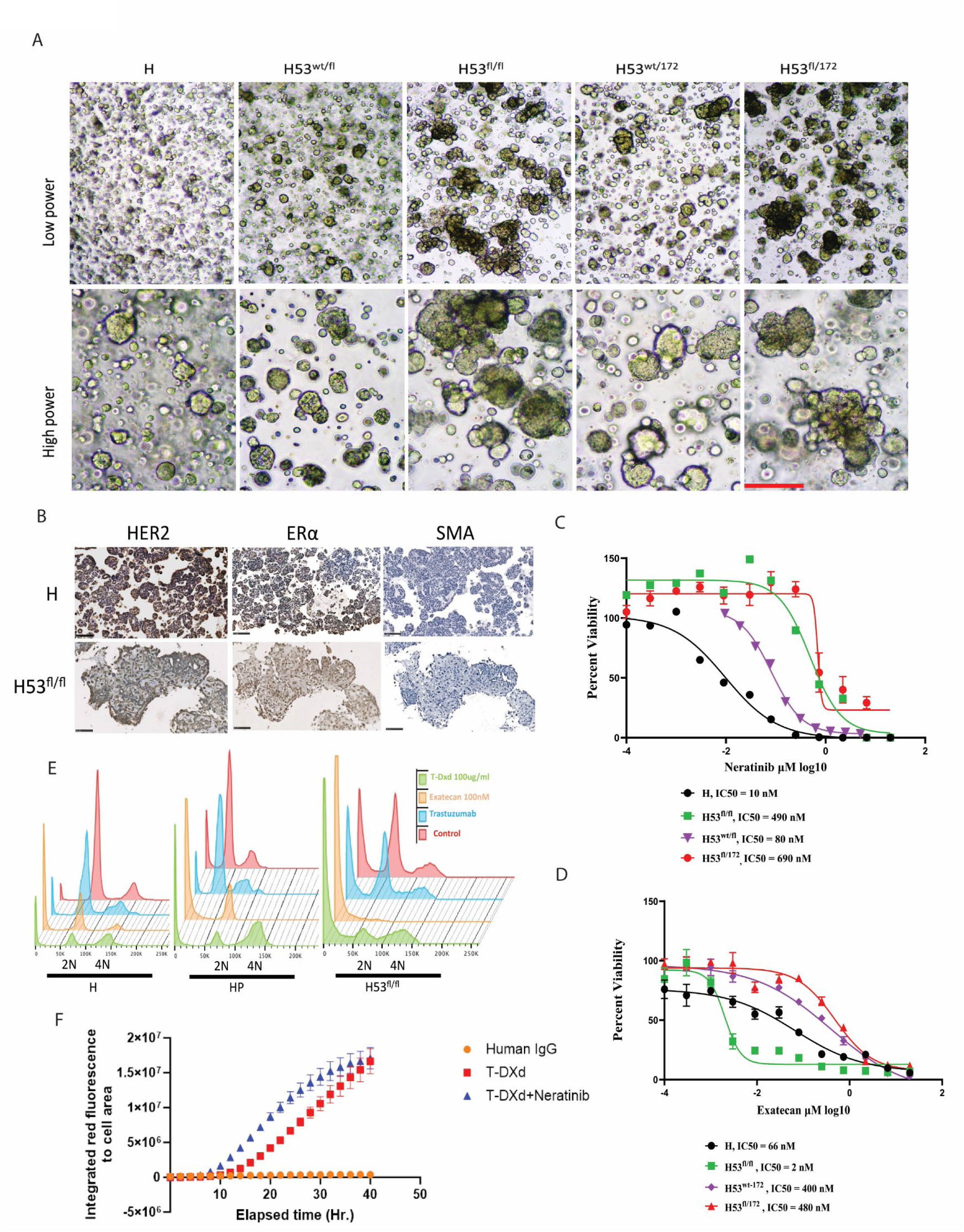
Organoids derived from H53 tumors show resistance to neratinib, but are more sensitive *in vitro* to TOP1 inhibitors. (A) Representative morphology of tumor organoids derived from mammary gland tumors from the H, H^wt/fl^, H^wt/172^, H53^fl/fl^, and H53^fl/172^ mice. (B) Immunohistochemistry (IHC) staining of HER2, ERα, and SMA images of tumor organoids from H and H53^fl/fl^ mice, respectively. Scale bars of low-power images are 5 mm. Scale bars of high-power images are 100 µm. (C) Representative IC50 assay of neratinib on organoids derived from mammary gland tumors from the H, Hwt/172, H53fl/fl, and H53fl/172 mice. (D) Representative IC50 assay of exatecan on organoids derived from mammary gland tumors from the H, Hwt/172, H53fl/fl, and H53fl/172 mice. (E) The cell cycle profile was assayed by FACS analysis with PI staining on the breast tumor organoids isolated from H, HP, and H53fl/fl mice without or with trastuzumab, exatecan, and T-DXd treatment. (F) Uptake assay in TD474 cell lines was performed using an Incucyte S3 Live Cell Analysis System (Sartorius).

We then examined the cell cycle effect of T-DXd, trastuzumab, and exatecan in H, HP, and H53^fl/fl^ tumor organoids. H and HP organoid cells showed similar cell cycle arrest patterns, while H53^fl/fl^ cells showed more dead cells in subG1phase when treated with T-DXd and exatecan (Fig. 2E). Additionally, we found neratinib can increase the uptake of T-DXd in HER2 positive breast cancer cells (Fig. 2F), hinting at the possible synergy of neratinib and T-DXd in treating these H53 tumors.

### The synergy of Neratinib and T-DXd in H53 tumors and their sensitivity to the TOP1 inhibitor are p53 transcription activity-dependent

To test our hypothesis that combination treatment is effective, we used a synergy assessment assay *ex vivo*. H53fl/172 and H53fl/fl tumor organoid cells showed sensitivity after 3 days of T-DXd single treatment. Nevertheless, we found very consistent evidence of drug synergy between neratinib plus T-DXd regardless of the H53 genotype (Fig. 3A). All three H53 genotypes showed synergy between neratinib and T-DXd. This is expected given that this drug synergy should occur at the level of receptor protein, HER2. We also detected mild synergy in HCI003, which is a PDX with both *HER2*, *PIK3CA*, and *TP53* mutations (Supplemental Fig. 4D).

**Fig. 3:**
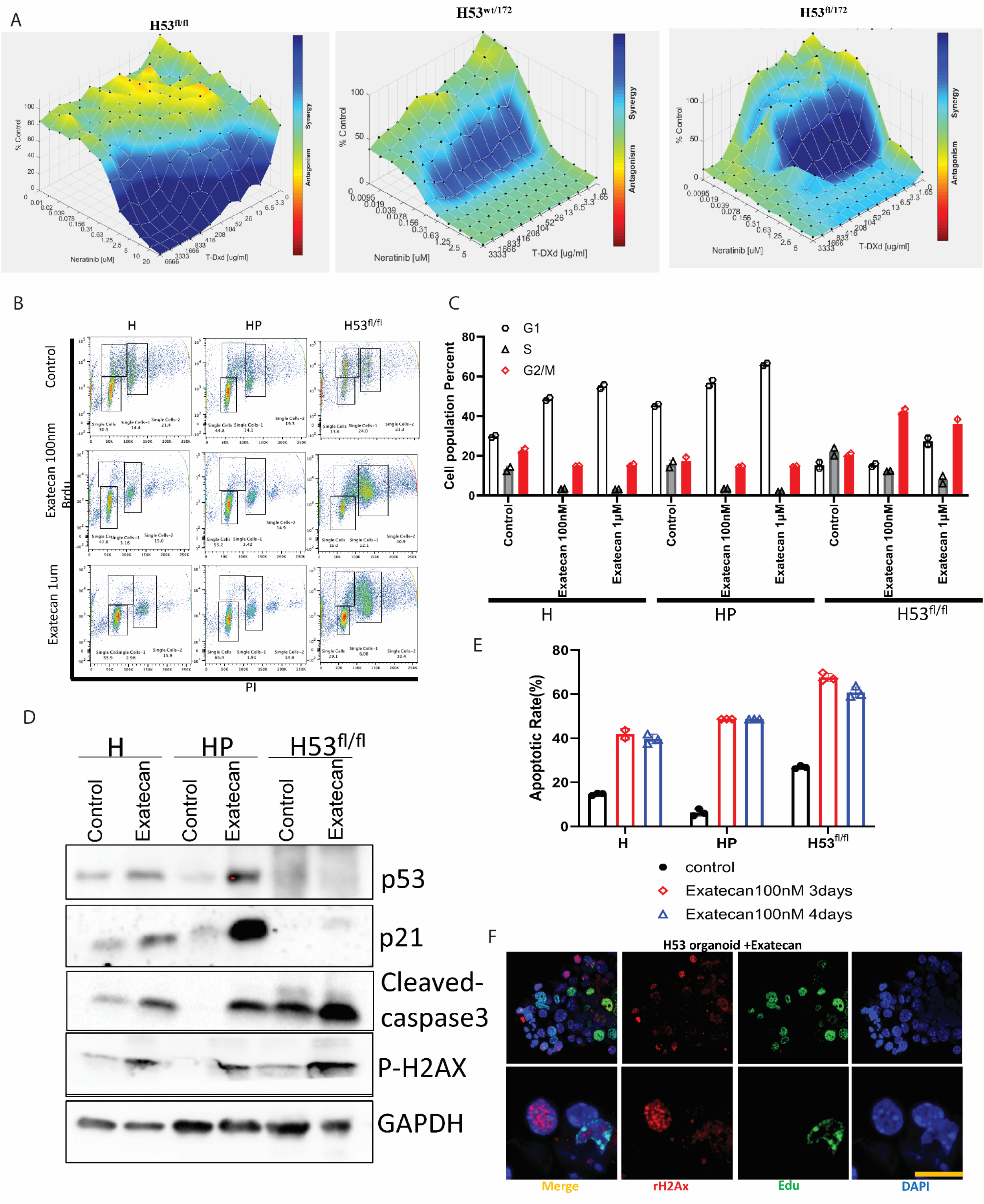
TOP1 inhibitor induce replication-associated DNA damage. (A) Testing the combined effect of T-DXd and neratinib in H53fl/fl, H53wt/172, and H53^fl/172^ organoid cell lines. Drug synergy, as per the Lowe model, is indicated by blue colors on the 3D surface, whereas green colors indicate additivity. (B) Cell cycle arrest assay using Flow cytometry assay with Brdu and Propidium iodide co-staining after 12 hours of treatment of exatecan at 100nM and 1uM on organoids derived from H and H53fl/fl mice. (C) The percentage of cells in each cell cycle phase was detected by Brdu-PI co-staining from tumor organoid cells isolated from H, HP, and H53^fl/fl^ mice. Error bars represent the standard deviation (SD) of triplicate technical replicates. (D) Western blotting p53, p21, phospho-CHK1, phospho-H2Ax, phospho-caspase3, and GAPDH from indicated mice breast-derived organoids after 2 days of exatecan treatment at 100nM. (E) Quantification of the percentage of apoptotic organoid cells. Apoptosis assay using FACS by annexin V staining after 3and 4 days treatment of 100nM exatecan on organoids derived from H, HP and H53fl/fl group mice. Error bars represent the standard deviation (SD) of triplicate technical replicates. (F) Immunofluorescence staining of γH2AX, EdU, and DAPI in 3D-grown organoids, as indicated. Error bar is 25µm.

To assess the drug sensitivity mechanism of exatecan in p53 and HER2 co-mutant organoid cells, we examined the cell cycle status using bromodeoxyuridine/fluorodeoxyuridine (abbreviated simply as BrdU) co-stained with propidium iodide, a fluorescent DNA intercalating dye detecting the incorporation of thymine analogs into newly synthesized DNA. This allows for the clear distinction of cells in G1 from the early S phase or the late S phase from G2/M. As shown in Fig. 3B-C, the H53 mutant tumor organoid cells arrest at the G2/M phase, while the H-only or HP tumor organoid cells arrest at the G1 phase when treated with exatecan. G2/M arrest continually accumulates the abnormal cancer cells and thus causes a mitotic catastrophe,^33^ which is a much more severe stress compared with G1 arrest. A apoptosis resulting from the abnormal cell cycle was also detected. More exatecan-induced cell death is observed in the H53 co-mutant tumor organoid cells (Fig.3 D-E). Western blotting results indicate that cell death is mediated by double-strand DNA-induced DNA damage as marked by γH2Ax and cleaved caspase3. The induction deficiency of p21, the direct transcription target of p53, in H53 co-mutant tumor organoid cells after exatecan treatment suggests this drug sensitivity may be p53 transcription activity-dependent. TOP1 cleaves single DNA strands to relax supercoiled DNA and alleviate the DNA helical constraints, while TOP2 cleaves double-stranded DNA followed by re-ligation. TOP inhibitors result in accumulating double-strand DNA breaks that mediate DNA damage, such as *PARP1* mutations, which need base excision repair. Meanwhile, free hydrolyzed ATP in the nucleus, which is usually used for TOP1 function, may cause chromatin remodeling-induced mitotic catastrophe when the processes of transcription, DNA replication, and DNA repair are stuck by TOP1 trapping. Our results here support these insights.

### *TP53* is the genetic determinant of drug sensitivity *in vivo*

We tested the impact of the p53 transgene on neratinib and T-DXd drug sensitivity. *In vivo*, drug testing of Neratinib plus T-DXd in mice is performed using a tumor organoid transplant model. Control arms include the two monotherapy arms (Neratinib alone and T-DXd alone) as well as a vehicle control arm. The drug dosage timelines are shown in Fig.4A. We conducted *in vivo* drug treatments on our organoid-transplanted mice. Neratinib, the antibody-drug conjugate, T-DXd, and the combination of trastuzumab deruxtecan and neratinib were each tested on H tumor organoid transplanted mice. *In vivo*, the antibody-drug conjugate T-DXd alone did not delay tumor formation or change the survival of H mice (Fig. 4B). Neratinib had the greatest effect on tumor formation in H, H53fl/172, and H53fl/fl tumor-burdened mice (Fig. 4B, Supplemental Fig. 3). T-DXd combined with Neratinib dramatically abrogated tumor formation over a 4-week period by two doses of 30mg/kg IV weekly (Fig. 4B). H, H53^fl/fl^, and H53^fl/172^ tumors showed different sensitivity responses to the Neratinib, T-DXd, and combination with Neratinib and T-DXd *in vivo* indicating the genetic determinates of the drug response mechanism (Fig. 4). Since neratinib is a pan HER inhibitor, H, H53^fl/fl^, and H53^fl/172^ tumors dependent on HER2 expression can be killed by neratinib. The sensitivity of these organoids to the ADC conjugate T-DXd was significantly different. Both *TP53*-containing samples, H53^fl/fl^ and H53^fl/172^, were sensitive to T-DXd (Fig. 4B), whereas H tumor organoids were sensitive to Neratinib and its combination with Neratinib (Fig. 4B-C). Neratinib has been shown to increase the internalization of trastuzumab-containing ADCs (Fig. 2F), which is the likely mechanism for this drug synergy. However, sensitizing the HER2; p53 co-mutant breast cancer to the T-DXd suggests that the dual targeting of HER2 by Neratinib and trastuzumab-based ADC is genetically dependent.

**Fig. 4:**
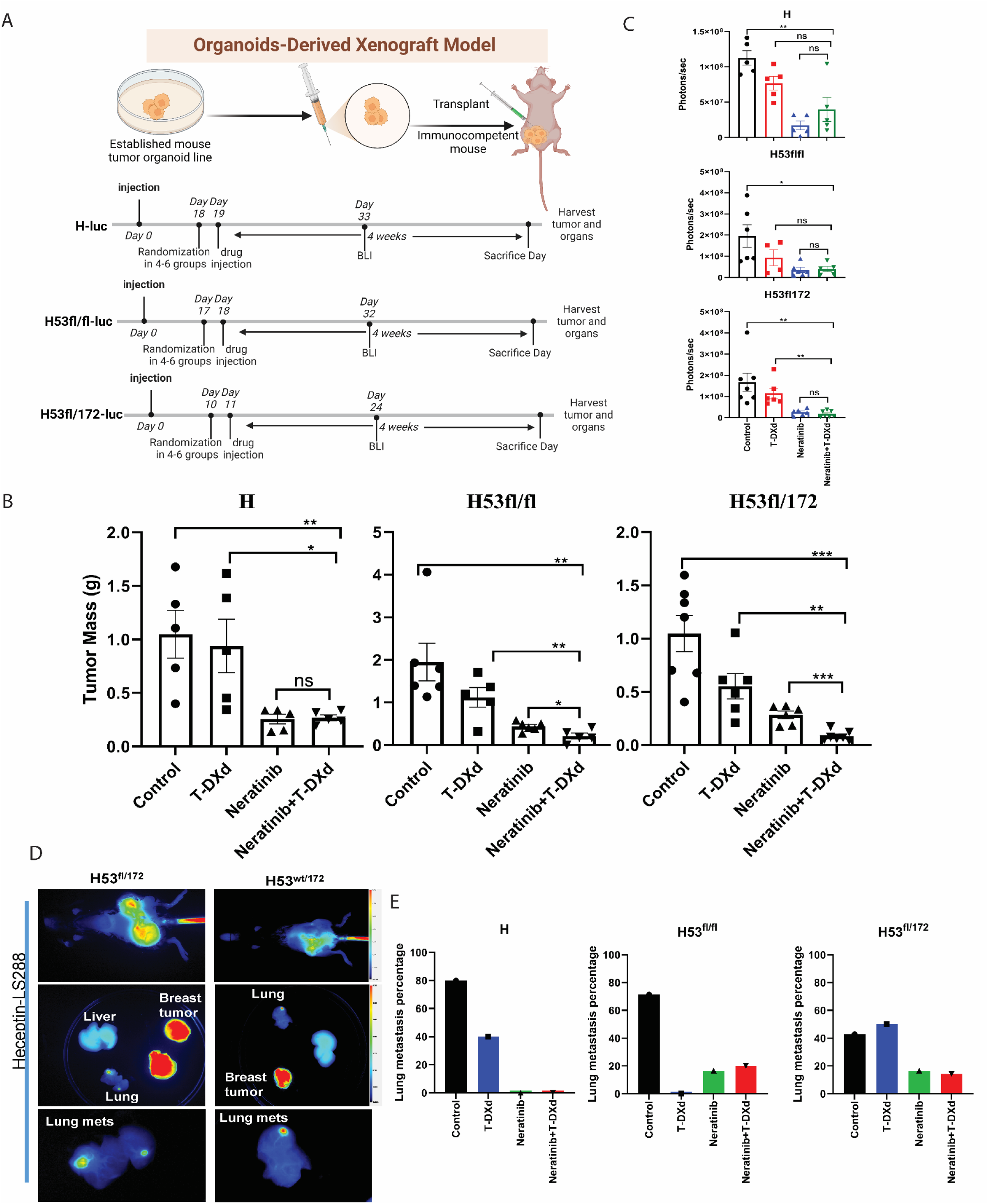
Drug testing *in vivo* on mice with T-DXd and Neratinib, which is a genetic determinant of drug sensitivity. (A) Schematic view of the drug treatment timelines in each transplanted model as indicated. Mice were treated with saline water, Neratinib only, T-DXd, and Neratinib plus T-DXd, respectively, for 4 weeks. Neratinib was given daily at 40mg/kg by oral gavage. T-DXd was given weekly at 30mg/kg by tail vein injection for two doses. (B) Tumor mass after treatment in transplanted models with different genotyped organoid cells. (C) Representative BLI graph with measurements of treated mice after harvesting. (D) Imaging of near-infrared fluorophore-labeled trastuzumab (NIR-trastuzumab) at 24 hours post-injection on the whole body of H53^wt/172^, H53^fl/172^ mice. (E) The percentage of lung metastasis in organoid transplanted mice of H, H53fl/fl, and H53fl/172. Data in (B) and (C) are plotted as mean ± SEM. ∗P < .05, ∗∗P < .01, ∗∗∗P < .001, and ∗∗∗∗P < .0001 as calculated by the Unpaired *t-*test.

Bioluminescence imaging (BLI) was performed serially on mice and is an accurate readout of tumor burden. The combination of neratinib and T-DXd showed the greatest tumor shrinkage as measured by both tumor mass and BLI (Fig. 4B-C). Tumor shrinkage with this combination was statistically significant compared to vehicle and either drug on their own, based on tumor mass measurements in H53fl/fl, and H53fl/172 mutant tumors.

To understand the p53 mutant’s contribution to metastases, we next examined the lung metastases in these *TP53* and HER2 co-mutant strains. The HER2^V777L^ mutation was shown to play a vital role in metastatic breast cancer. As shown in Fig. 4D, H53wt/172 and H53fl/172 show lung metastasis in the adenovirus-based Cre-generated mice. To test spontaneous metastasis status in H, H53fl/fl, and H53fl/172 tumors, these tumor organoid transplanted mice were examined for metastases in other organs like the lung or liver. Lung metastases were seen in both H, H53null, and H53null/172 tumor organoid transplanted mice (Fig. 4E, Supplemental Fig. 4A-B). Clinically, slowing growth of the primary tumor leads to high percent metastasis. In H53wt/172 tumor organoid transplanted mice, lung metastases were seen at eight months after adenovirus Cre injection into the mammary glands (Fig. 4D). Lung metastases were observed in more than 50% of transgenic H53^fl/fl^ mice (Fig. 4E). To improve visualization of both primary tumor and metastases, we conjugated trastuzumab with the near-infrared (NIR) fluorophore, LS288. H, H53^fl/fl^, and H53^fl/172^ tumor-bearing mice injected with this NIR-trastuzumab imaging agent showed tumor-specific uptake at 24 hours in the breast and lung but not in the liver or spleen (Supplemental Fig. 2A-B, Fig. 4D-4E). The imaging data suggested that the lung is a metastatic site for the HER2^V777L^-expressing breast tumor cells in our mouse models. Therefore, we conclude that HER2 is the dominant genetic determinant for lung metastasis, and p53 may have a role contribution to lung metastasis.

Metastases, rather than growth of the primary tumor, are the cause of death in the vast majority of breast cancer patients. Therefore, we tested the ability of Neratinib plus trastuzumab deruxtecan and a single treatment of each of these two drugs individually to treat lung metastases in H, H53^fl/fl^ and H53^fl/172^ tumor burdened mice (Fig. 4E). Female mice were orthotopically injected with luciferase-labeled H, H53^fl/fl^ and H53^fl/172^ breast cancer organoids into the fourth mammary gland, located in the inguinal region of the mouse. After 11-19 days, drug treatment with vehicle, neratinib, T-DXd, and neratinib plus T-DXd was started. After 4 weeks of drug treatment, mice were euthanized, and bioluminescence imaging and histology were performed on the dissected lungs. As shown in Supplemental Fig. 3A, the lungs of mice treated with Neratinib and Neratinib plus T-DXd showed a significant decrease in luciferase signal compared to the vehicle control. Similarly, tumor mass measurements (Supplemental Fig. 3B) showed the effects of drug treatment on the primary tumor in the fourth mammary gland. Histologic examination of the lungs showed numerous lung metastases in the vehicle group and almost no lung metastases in both drug treatment groups (Supplemental Fig.4A-B). These data represent pre-clinical evidence that Neratinib plus T-DXd are effective for treating HER2 and p53 co-mutated metastatic breast cancer. As expected, the prolonged survival provided time for the development of lung metastasis in the H53 mutant organoid transplanted mice, suggesting that our organoid-derived xenograft model can be used to study the mechanisms and therapy of breast cancer metastasis.

### *TP53* mutation sensitizes the cells to the TOP1 inhibitor in a transcription activity-dependent way

To further elucidate the sensitivity mechanism of tumors of H53 mutant to the TOP1 inhibitor, we performed cell synchronization using double thymidine treatment in both H only and H53 co-mutant tumor organoid cells. Results showed cell cycle arrest in S phase after 20 hours of exatecan treatment in H53 co-mutant tumor organoid cells, while there is an obvious G2/M arrest at the 24-hour treatment timepoint (Fig. 5A). The rewiring signaling was examined using the same cells. P-CHK1 was upregulated at the earliest response to γH2Ax as expected in H53 co-mutant tumor cells, then causing the cleaved-caspase-mediated cell death (Fig. 5B). The P-CHK1 signal showed an obvious increase along with the cell cycle arrest release. This suggests CHK1 played different roles in DNA damage-induced cell cycle arrest in human breast cancer^34^.

**Fig. 5:**
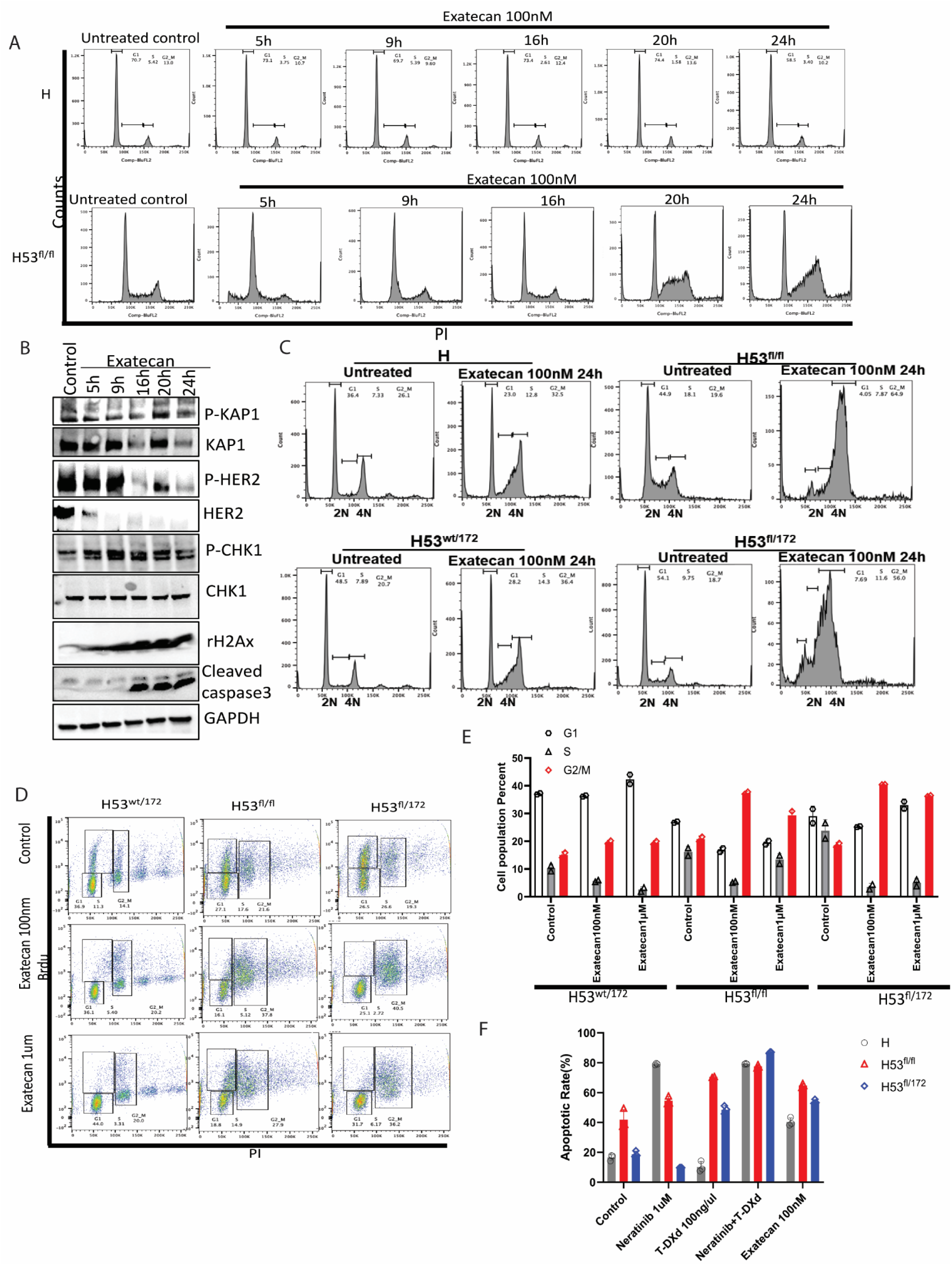
The genetic determinant of drug sensitivity is p53 transcription activity dependent. (A) Cell cycle assay using flow cytometry with Propidium iodide (PI) staining after treatment of exatecan at 100nM at indicated timepoint on organoids derived from H (Top) and H53fl/fl (bottom) mice. (B) Western blotting of KAP1, p21, phosphor-KAP1, phospho-CHK1, total CHK1, phospho-CHK2, total CHK2, phospho-HER2, total HER2, phospho-H2Ax, phospho-caspase3 and GAPDH from exatecan-treated H53 null organoid cells in Figure 3A. (C) The cell cycle profile was assayed by FACS analysis with PI staining on the breast tumor organoids isolated from H53wt/fl, H53fl/fl, and H53fl/172 mice. (D) The percentage of cells in each cell cycle phases was detected by BrdU-PI co-staining from tumor organoids cells isolated from Hwt/172, H53fl/fl, and H53fl/172 mice. Error bars are based on triplicated technical replicates. (E) Cell cycle phase analysis based on the FACS result from figure 5D. Error bars are based on triplicate of technical repeats. (F) Quantification of the percentage of apoptotic organoid cells. Apoptosis assay using FACS by annexin V staining after 3 days of treatment of 100nM exatecan on organoids derived from H, HP, and H53fl/fl group mice. Error bars are based on triplicate of technical repeats.

Cell cycle status of H only, H53^wt/172^, H53^fl/172^, and H53fl/fl were compared in the exatecan treatment scenario. Both H53fl/fl and the H53fl/172 tumor organoid cells showed cell cycle arrest (Fig. 5C-E). We confirmed this G2/M arrest using BrdU staining in the H53 co-mutant tumor organoid lines (Fig. 5D-E). Compared with *TP53* deletion, the R172H mutation of *TP53* blocks the p21 expression and phenocopies the G2/M arrest upon TOP1 inhibition, suggesting the transcription activity of *TP53* drives the sensitivity response. The combination of T-DXd and Neratinib shows an effective result when treated with *TP53*-mutated organoids (Fig. 5F).

### Histone modification proteomics using organoids derived from H and H53^wt/fl^, H53^wt/172^, H53^fl/fl^, and H53 ^fl/172^ mice tumors

As T-DXd treatment targets TOP1 and its related DNA machinery in DNA replication, DNA repair complex, and transcription clusters, we examined the chromatin status of the HER2^V777L^ and H53 co-mutant tumor organoid cells with respect to drug sensitivity. As transcription is the predominant response to histone modification, we examined chromatin status by checking the histone modification in the HER2 mutant-only and H53 co-mutant tumor organoid cells.

Mass spectrometry was used to detect histone modifications in HER2 mutant-only and H53 co-mutant tumor organoid cells. PCA analysis in Fig. 6A demonstrates a difference in H53 co-mutant organoid groups versus the HER2 mutant-only groups, indicating that histone modifications may play a previously unrecognized role in sensitizing tumors to treatments. Striking upregulation of histone acetylation was identified in H53 co-mutant organoid groups compared with the H-only group (Fig. 6B). Since histone acetylation neutralizes the histone octamer to dissociate DNA, DNA with histone acetylation is in a more open state and more accessible. More transcription factors can bind to these open areas in response to accelerated tumorigenesis and reduced survival signal. Meanwhile, the open status of DNA sensitizes these cells to the DNA-targeting drugs like TOP1-DNA covalent or DNA damage-targeted drugs. These offer a novel angle and new insight for future drug design and therapeutic strategies. The Venn diagram in Fig. 6C shows there are more shared histone modifications than are unique to each group. The top four shared modifications among these H53 co-mutant organoid groups are H3K9ac, H2AzK4acK7ac, H2A3K13ac, and H3.3K36me1 (Fig. 6C). The modification peaks are consistent among each organoid line (Fig. 6D). We further confirmed the histone acetylation of H3K9 upregulation in the H53 co-mutant organoid groups via western blotting (Fig. 6E). H3K9ac is one of the main histones marks for active promoters to turn on transcription. We also examined the TOP1, P-HER2, HER2, KAP1, r-H2Ax, p53 and p73 level in all H53 mutant cells. *TP53* mutation upregulates the H3K9ac, which can be a compensatory feedback regulator. However, as a target of H3K9ac, TOP1’s instability is related to protein interaction with mutated p53 ^35^ and TOP1 inhibitor-induced degradation. ATM-based signaling is distinct between *TP53* loss of function and gain of function, suggested by the difference in KAP1 level (Fig. 6E). DSB of DNA damage-inducing cell death may be mediated by p73 in the p53 deletion and mutated scenario (Fig. 6E).

**Fig. 6:**
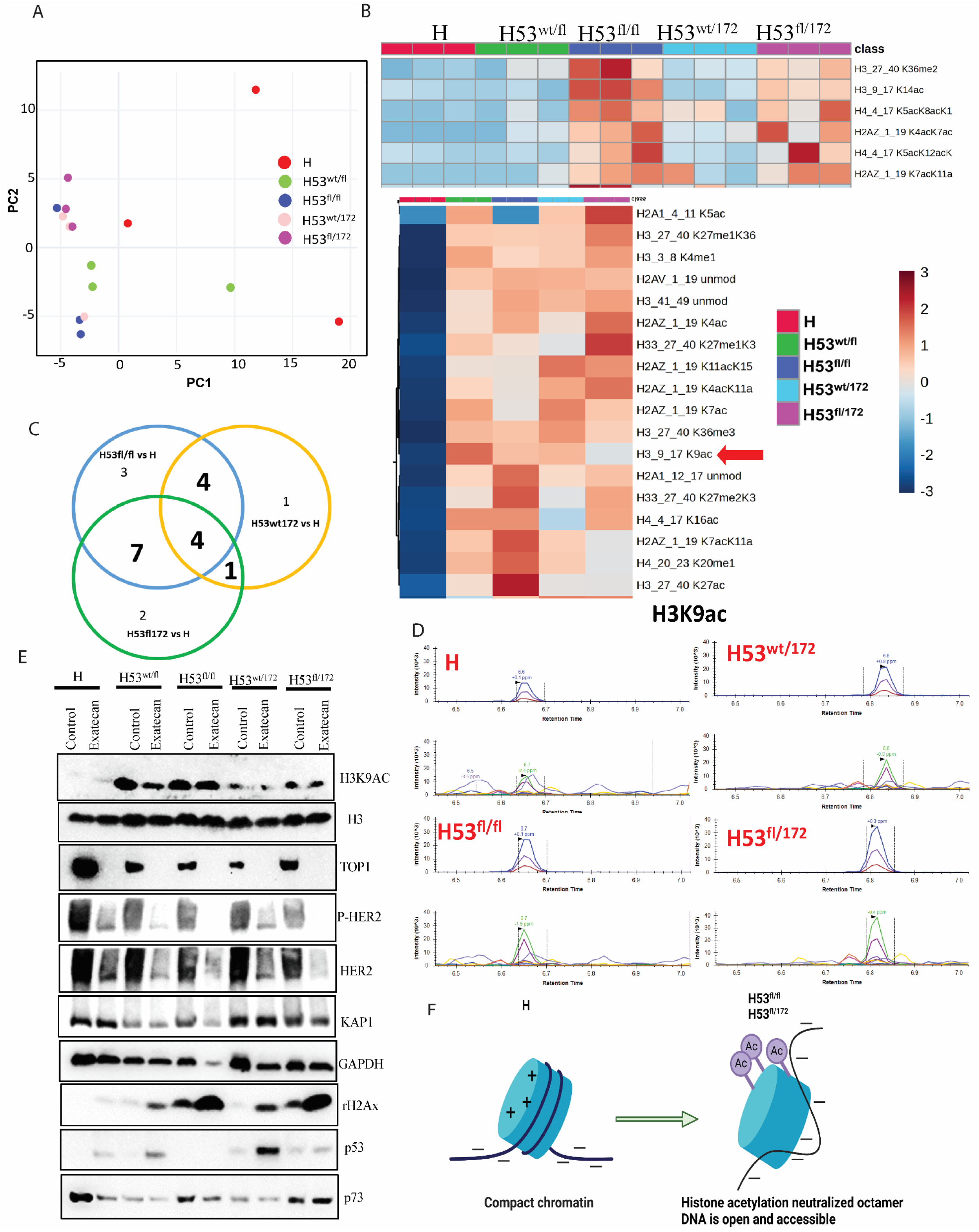
Histone modification proteomics using organoids derived from H and H53wt/fl, H53wt/172, H53 fl/fl, and H53 fl/172 mice tumors. (A) PCA analysis confirms transgene-based lineage relationship and genetic heterogeneity in histone samples. (B) Heatmap showing the top 25 histone modifications of DEGs (abs(log_2_ FC) > 1, P_adj_ < 0.05) between H, H53 co-mutated organoid samples. (C) Venn diagram shows the shared histone modification marks among H53 co-mutated organoid samples compared with H-only groups. (D) The histone modification identified peaks in the mass spectrometry system. (E) Western blotting results detecting the H3K9ac, H3, TOP1, p-HER2, HER2, KAP1, GAPDH, γ H2Ax, p53, and p73 level in H, H53 co-mutated organoid samples. (F) The model shows histone acetylation in chromatin remodeling by neutralizing the histone charge.

Histone acetylation can neutralize the octamer to loosen the binding of DNA, thus turning on gene expression (Fig. 6F). H3K9ac and H3K14ac are predictors of active promoter status. As in aggressive tumor cells, accelerated cell growth calls for an overactive transcription process for nutrition and microenvironmental support. Additionally, transcription is reported as a therapy vulnerability for several drugs. Our findings explain the drug-sensitive mechanism of p53 mutant cells’ response to the T-DXd. These findings also confirm chromatin remodeling-related activities like transcription can be a therapy vulnerability. Combined, these findings open a new window for future therapeutic strategies for p53 mutated cancer types.

### H3K9AC marks the promoters of p53 and TOP1 in all breast cancer types. Combined Neratinib with T-DXd shows synergy in PDX

Using Chip-seq data, we found that H3K9ac marks the promoters of *TP53*, *p73* (but not p63), *TOP1, TOP2A, TOP2B, ATM, KAP1*, and *NPAT* in all breast cancer types. We also mapped the levels of both H3K9ac and H3K9me3 over these genes in several breast cancer cell lines using ChIP-seq data available on GEO (GSE85158). The cell lines examined include the five major breast cancer subtypes, including two ER-positive subtypes (Luminal-A and Luminal-B), the HER2-positive subtype, and two triple-negative subtypes (TNBC-Basal and TNBC-Claudin Low). Additionally, normal-like immortal breast cells were included in the analysis. Data displayed are normalized counts. H3K9ac levels over the *TP53* gene in the different breast cancer cell lines and normal-like cells follow a broadly similar pattern with some variations in their magnitude. Regarding the HER2 positive subtypes, HCC1954 shows relatively high H3K9ac levels, particularly in replicate 1 (Fig. 7A). The levels of H3K9ac over TOP1 using the same breast cancer cell lines from GEO (GSE85158) were also obtained (Fig.7A). Correspondingly, H3K9me3, which is the mutually exclusive hallmark of H3K9ac, shows no specific enrichment at the promoters of these genes throughout all the breast cancer cell types (Fig.7B).

**Fig. 7:**
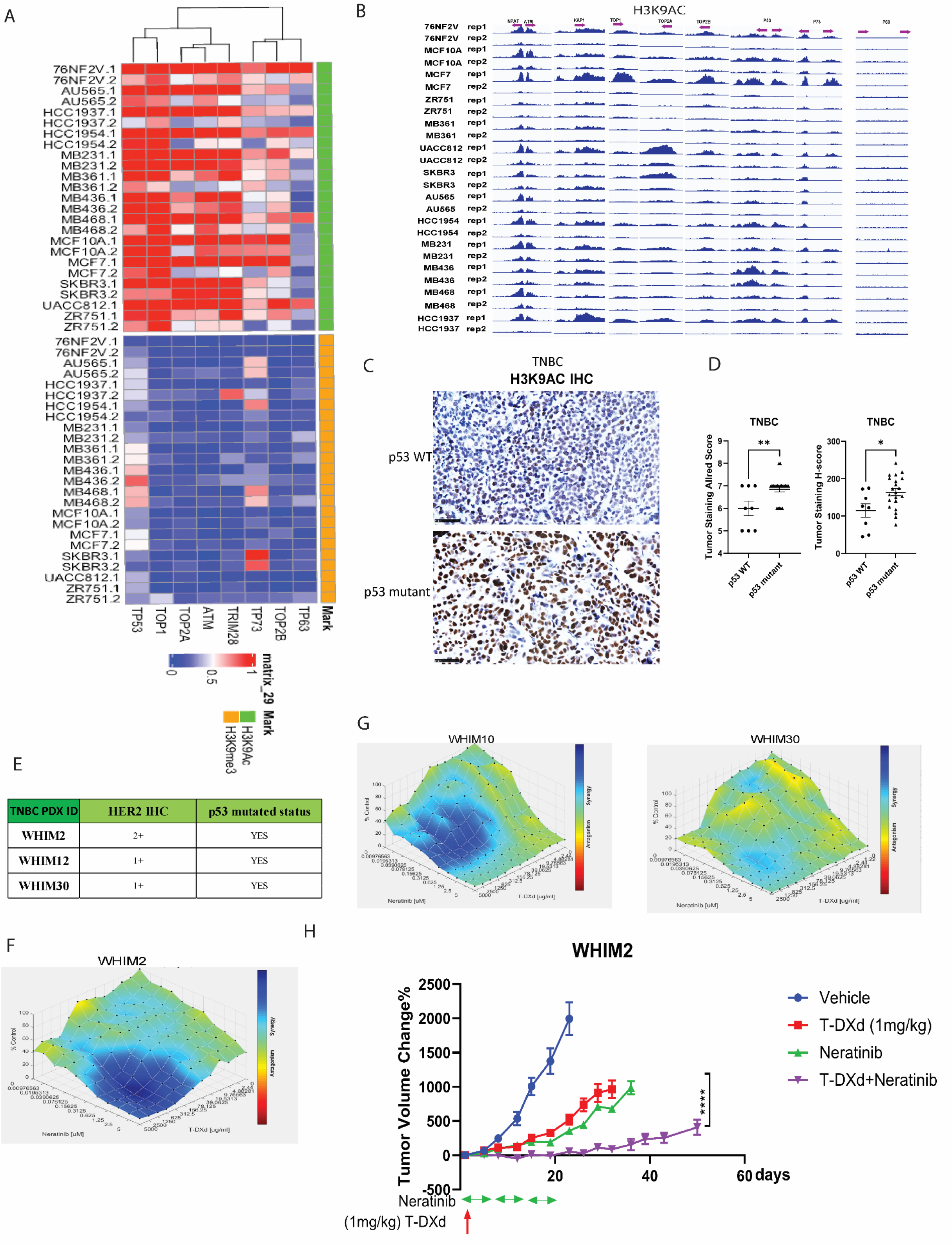
H3K9AC marks the promoters of P53 and TOP1 in all breast cancer types. Human PDX HCI003 shows mild synergy when combined with neratinib and T-DXd. (A-B) H3K9AC marks the promoters of p53, p73, ATM, KAP1, TOP1, TOP2A, TOP2B, and NPAT, but not p63 in all breast cancer types. H3K9me3 doesn’t mark the p53, p73, p63, ATM, KAP1, TOP1, TOP2A, TOP2B, and NPAT promoters in all breast cancer types. (C)The PDX tumor microarray of triple negative breast cancer showed increased H3K9AC IHC signal. (D)Tumor staining Allred score and H score of H3K9AC IHC signal from tumor microarray in Fig 7C. (E)The information of PDXO is in the TNBC PDX list. (F-G) Synergy of neratinib and T-DXd in human PDX of WHIM2, WHIM12, and WHIM30. (H) The in vivo treatment of the WHIM2 with neratinib and T-DXd.

H3K9AC expression in both wild-type and p53-mutated PDX was validated using a tumor tissue microarray (Table 2). Consistent with our histone mass spectrometry data in Fig. 6, there is a significant upregulation of H3K9ac in triple negative breast cancer (TNBC) with p53 mutations (Fig.7C). Both tumor staining Allred score and H-score showed a significant upregulation of H3K9ac in TNBC (Fig.7D).

To further confirm the mechanism of drug sensitivity in H53 mutant breast cancer, we tested the exatecan on several TNBC cell lines with *TP53* mutations. We found that there is a strong synergistic effect when combined with neratinib and T-DXd in PDX organoid model of TNBC with HER2 positive expression (Fig. 7E-G). Similar to the results of H53 mutant breast cancer organoids (Fig 4), the *in vivo* treatment on these TNBC cell lines showed similar sensitivity to T-DXd (Fig. 7H), suggesting that T-DXd can benefit patients with TNBC. T-DXd plus Neratinib can be an effective clinical treatment for patients with HER2 low TNBC breast cancers and such treatment has been clinically practiced for the past two years.^36^ In conclusion, these data suggest that T-DXd can be used as a treatment strategy for tumors containing *TP53* mutations.

### The mechanism working model

The mechanism of drug sensitivity is summarized in the working model. In HER2 and *TP53* co-mutated cells, increased histone acetylation loosens histones from DNA, making it more accessible to transcription factors and DNA-targeted drugs. Upon T-DXd treatment, exatecan forms a TOP1-exatecan covalent complex during DNA replication, causing both DSBs and SSBs. This DNA damage activates the ATM-KAP1-pCHK1-p53 pathway for repair. Normally, p53 induces p21 to regulate cell cycle progression during repair. However, in H53 mutant tumor organoids, defective p53 transcription impairs this response, causing cells to arrest at G2/M phase and undergo cell death.

## Discussion

In this study, we demonstrate that targeting topoisomerase I (TOP1) is an effective therapeutic approach for *TP53*-mutant breast cancers, especially in the context of HER2 co-mutations. Our findings reveal that *TP53*-mutant tumor organoids are highly sensitive to the TOP1 inhibitor exatecan and to the HER2-directed ADC trastuzumab deruxtecan (T-DXd). Notably, combining T-DXd with the HER2 inhibitor neratinib enhanced ADC uptake and produced a synergistic cytotoxic effect across various *TP53* genotypes. Mechanistically, *TP53*-deficient cells exhibited S/G2 phase arrest and impaired p21 induction, suggesting transcriptional control as a critical determinant of drug response. We also observed elevated histone acetylation in *HER2/TP53* co-mutant organoids, indicating chromatin regulation as a potential vulnerability. These insights highlight the therapeutic potential of combining HER2-targeted therapies with DNA-damaging agents and chromatin-modifying drugs in genomically unstable tumors.

Despite these promising findings, our study has several limitations. First, while we used genetically defined organoid models and mouse tumor systems, these models do not fully recapitulate the complexity of human tumor microenvironments and immune interactions. Second, although *TP53* mutations were associated with differential cell cycle arrest and drug sensitivity, the specific contributions of gain-of-function versus loss-of-function *TP53* mutations warrant further exploration. Mitotic arrest^37^ and ATM loss sensitizes tumor cells.^38^ Studies indicate that *TP53* loss-of-function (LOF), rather than gain-of-function (GOF), drives tumor maintenance in many cancers, ^39^ while in colorectal cancer, combined LOF and GOF mutations enhance metastasis but exert distinct tumorigenic roles.^40^ Third, our data primarily focus on *HER2/TP53* co-mutant breast cancer, and additional studies are needed to assess whether these findings are applicable to other *TP53*-mutant cancers, such as triple-negative breast cancer or gynecologic malignancies. *TP53* mutations occur in 50% of all cancers and up to 90% of ovarian and uterine cancers. ^41^ This mechanism could extend as a therapeutic insight in triple-negative breast cancer (TNBC). Another possible mechanism involves R-loops and mitotic catastrophe. While TOP1 chelate-induced DSBs mainly arise from DNA replication, they can also result from accumulated positive supercoils.^42^ Without TOP1 relieving this tension, positive supercoils ahead of the replication fork hinder progression, causing fork collapse and lethal DNA lesions that trigger cell death.

Looking forward, future studies should validate these findings in patient-derived xenografts (PDX) and clinical biopsy samples, particularly in the context of ongoing clinical trials combining T-DXd with neratinib (NCT05372614). Additionally, it will be important to explore the therapeutic potential of combining TOP1 inhibitors with inhibitors of DNA repair (e.g., Tdp1) or cell cycle checkpoints (e.g., ATM, Chk1/Chk2). Given the transcriptional and chromatin remodeling dependencies identified in this study, epigenetic therapies such as HDAC inhibitors may further enhance treatment efficacy and help overcome resistance mechanisms. Integrating multi-omics profiling may also help stratify patients based on chromatin states or DNA repair deficiencies to guide precision therapy.

In conclusion, our study provides mechanistic insight and preclinical evidence supporting the use of HER2-targeted ADCs and TOP1 inhibitors in HER2;*TP53* co-mutant breast cancers. The observed synergy with neratinib, along with chromatin-based vulnerabilities, opens new avenues for combination therapies tailored to the genetic and epigenetic landscape of tumors. These findings have the potential to inform future clinical trial design and extend therapeutic strategies to other *TP53*-mutant cancers characterized by replication stress, chromatin remodeling, and resistance to conventional treatments.

## Acknowledgements

We would like to express our deepest gratitude to our colleagues for their invaluable guidance, support, and encouragement throughout this project. I also thank the patient advocates and members of the research community who have inspired and informed my work. Special thanks to Dr. Thomas Walsh, Dr. Jason Weber, and Dr. Katherine N. Weilbaecher for their resource sharing, thoughtful feedback, and collaboration. We would like to thank Siteman Cancer Center’s scientific editor, Megan Noonan, PhD, for her expertise and insightful suggestions, which enhanced the clarity and quality of this work.

## Materials and Methods

### Mice

HER2^V777L^ transgenic mice were generated using TALEN-based genome editing as previously reported.^32^ *P53^R172H^* mice were shared from Frederick National Laboratory, strain #:01XAF. *P53^fl/fl^*mice were purchased from the Jackson Laboratory, strain #:008462. All animal studies were approved by the Institutional Animal Care and Use Committee (IACUC) at Washington University in St. Louis.

### Induction of Cre-mediated recombination

To study the contribution of co-mutation of HER2 and *TP53*, HER2^V777L^ mice were crossed with *TP53^fl/fl^* or *TP53^R127H^* mice to generate H53 mice. All experiments were done using congenic mice (C57BL/6J mice) and all experiments were performed with littermate control. For adenovirus Cre-mediated recombination, 8-10-week-old mice were injected orthotopically into the mammary fat pad with adenovirus Ad5CMVCre, which was purchased from the Viral Vector Core Facility of the University of Iowa, College of Medicine (titer range 1×10^10 to 7×10^10 pfu/ml).

### *Ex vivo* single drug sensitivity assay

Organoids were removed from Matrigel using Cell Recovery Solution (Corning), washed, and resuspended in media with 6% Matrigel (GFR). 25ul of organoids in 6% Matrigel were added to a 384-well black wall, clear bottom, and tissue culture plate and incubated overnight at 37 degrees C, 5% CO2. DMSO-based drugs were serially diluted 3-fold in 100% DMSO and then diluted into mouse breast tumor media in a 96 well plate at 2X the final desired concentration. Aqueous drug serial dilutions were made directly in media in the 96-deep well plate. The drugs, diluted in mice tumor media, were tested in duplicate with 25ul of drug added to the cells. Control wells were also prepared and added with a vehicle in the media to serve as 100% viability controls. Wells without cells serve as 0% viability controls. Organoids and drugs were incubated for six days. Viability was measured by adding 25ul of Cell Titer Glo 3D (Promega) to each well. Sealed plates were shaken for 5 minutes and incubated for another 30 to 60 minutes at room temperature (signal stable for ∼4 hours). Luminescence was measured with a Tecan plate reader.

### *Ex vivo* drug combination assay

Organoids were removed from Matrigel with Cell Recovery Solution and digested with TrypLE. After washing with Ad DME/F12 media, cells were diluted into cold mouse breast tumor media plus 6% Matrigel (GFR) and 20ul were added per well of a 384 well black wall, clear bottom, tissue culture-graded plate. The plate was incubated at 37 degrees C and 5% CO2 overnight. DMSO-based drugs were first serially diluted in 100% DMSO and then diluted into mouse breast tumor media in a 96-deep well plate. Aqueous drug serial dilutions were made directly in media in the 96-deep well plate. The two drugs to be compared were prepared at 3X the final desired concentration and the serial dilutions were 2-fold. Using the adjustable spacing Integra Voyager pipettor, 20ul of each of the two drugs were transferred from the 96 well intermediate drug dilution plate and added to the 384 well assay plate in a grid format as described in Griner, et al.^43^ Each drug was also assayed as a single drug with 20ul media making up for the lack of a 2nd drug. Additional wells were seeded with cells as vehicle controls for the 100% viability and wells without any cells for the 0% viability controls. Organoids and drugs were incubated for six days. Viability was measured by adding 30ul of Cell Titer Glo 3D (Promega) to each well and incubating as described. The percent viability for both assays was determined by first calculating the average vehicle value and the average “no cell” value. Then, percent viability was determined with the formula: Percent viability equals 100 minuses (ave viability control minus vehicle value)/(ave viability control minus ave no-cell control) X 100. IC50 values and graphs were calculated using GraphPad Prism 5 for Windows. The Synergy mapped to D-R (Loewe) plots were prepared using the freeware program Combenefit 2.021 for Windows from the University of Cambridge, Cancer Research UK.

### Luciferase labeling

Organoid cells were transduced with a lenti-LUC luciferase reporter. Cloning selection was performed to isolate stable clones with luciferase expression. Cell lines with labeling were confirmed using in vitro bioluminescence signal quantification.

### In vivo therapy

Drug treatments were conducted on three different H53 mutant cancer models. 10-12-week-old female H53 mutant mice with orthotopic Adenovirus-Cre injection on the mammary gland, H53 mutated organoid transplanted mice, and HCI003 PDX mice. Female H53 mutant mice at 18-32 weeks old were randomized to 4-6 treatment groups (n=5-7 per group): 1) vehicle, 2) neratinib, 3) T-DXd only, and 4) combined neratinib and T-DXd. Mice were subjected to the treatment from the 11^th^, 18^th^, or 19^th^ -day post-injection. This drug combination was chosen because we have already determined that H53 mutant mouse breast tumors are neratinib resistant, but sensitive to T-DXd.

Neratinib was dosed at 40 mg/kg daily by oral gavage to animals. T-DXd was dosed at 30 mg/kg weekly by tail vein injection. All animal experiments were conducted in compliance with federal guidelines and approved by the Institutional Animal Care and Use Committee at Washington University School of Medicine.

### In vitro therapy

*In vitro* experiments: H53 mutant breast cancer organoids and PDX HCI003 human breast cancer organoids were routinely cultured in 3D culture with Matrigel as previously described.^44^ Drug concentration added as indicated.

### Immunohistochemistry and Immunofluorescence of paraffin-embedded sections

Tumor tissues were fixed with 10% formalin, embedded in paraffin, and cut into 5-µm sections (Digestive Diseases Research Core, Washington University School of Medicine). Sections were de-paraffinized, hydrated, and treated with heat-activated antigen unmasking solution (Vector Laboratories). Immunostaining was performed using the antibodies listed below. For immunofluorescence staining, antibody binding was visualized with Alexa Fluor 488 or 555 or 647 fluorochromes, then counterstained with DAPI-containing mounting media (Sigma). For immunohistochemistry staining, DAB substrate (Cell Signaling) was applied, then counterstained with hematoxylin (Thermo Fisher Scientific). Primary antibodies used in the study include rabbit ERBB2 (CST, 2165S), mouse ERBB2 (CST, AF2967-SP), Smooth Muscle Actin Antibody (Santa Cruz Biotechnology, SC-53142), ER***α*** Antibody (D-12) (Santa Cruz Biotechnology, SC-8005), GATA-3 Antibody (Santa Cruz Biotechnology, SC-268), Secondary antibodies used included anti-rabbit IgG (H+L) Alexa Fluor 555 (Invitrogen, A-31572), anti-goat IgG (H+L) Alexa Fluor Plus 488 (Invitrogen, A32814), Goat anti guinea pig (H + L) FITC (Fitzgerald Industry International, 43R-1095), SignalStain Boost IHC Detection Reagent (HRP, Rabbit) (Cell signaling technology, 8114S).

### Establishment of breast organoid

Isolation of epithelial organoids from the mammary gland tumor from HER2^V777L^ only, HER2^V777L^; *P53^WT/R172H^*, HER2^V777L^; *P53^null/R172H^*, HER2^V777L^; *P53^wt/null^*, HER2^V777L^; *P53^null^* transgenic mice as previously described.^4^ Briefly, isolated organoids were embedded in Matrigel (BD Biosciences) and seeded in 6-well plates. The cells were overlaid with 2 mL/well basal culture medium supplemented with penicillin/streptomycin, 10 mmol/L HEPES, Glutamax, 1 × B27 [all from Thermo Fisher Scientific], 125 µM N-acetyl-cysteine (Sigma), 50 ng/ml murine epidermal growth factor (EGF, Invitrogen), 10% Rspo1-Noggin-conditioned medium.

### Organoid transplant mouse model

Organoid cells were collected and digested with TripLE for 5-10 minutes. The cells were counted, and 1 -2 x 10^6^ organoid cells were injected into the fourth mammary fat pad of congenic mice. On the fourth day, tumors were treated with a drug and collected after 4 weeks post-treatment.

### H53 mutant mouse Lung Metastasis Treatment

Luciferase-labeled H53 mutant breast tumor organoid cells (5 × 10^5^) were orthotopically injected into the inguinal mammary gland of female mice. On days 11, 18, or 19, when treatment was initiated with saline water, neratinib only, T-DXd, or neratinib plus T-DXd, respectively, for 4 weeks. Neratinib was given daily at 40mg/kg by oral gavage. T-DXd was given weekly at 30mg/kg by tail vein injection for two weeks.

### *In vivo* bioluminescence imaging (BLI)

For BLI of live animals in the Washington University Molecular Imaging Center (MIC), mice were injected intraperitoneally with 150 μg /g D-luciferin (Gold Biotechnology, St. Louis, MO) in PBS, anesthetized with 2.5% isoflurane, and imaged with a charge-coupled device (CCD) camera-based system (IVIS 50, PerkinElmer, Waltham, MA; Living Image 4.3.1, exposure time 10-60 seconds, binning 8, field of view 12cm, f/stop 1, open filter, ventral view). Regions of interest (ROI) were defined manually over the lower abdomen using Living Image 2.6 with measurements reported as photons/sec.

### Pearl Image

NIR-trastuzumab was injected into the tail vein of H or HP organoid transplanted mice bearing breast tumors one day prior to imaging. The near-infrared imaging was taken using the Pearl Trilogy (LI-COR, Lincoln, NE) in the Washington University Molecular Imaging Center (MIC). The PearlCam and ImageStation software was used for the measurement and analysis.

### Histone Extraction

Histone extraction method was adapted from Homsi et al^45^. Tumor cell pellets from respective conditions were solubilized in 600 µl of acidified 80% acetonitrile (0.25 M HCL) and lysed by probe sonication. The lysate was centrifuged at 15,000 rpm for 2 minutes, and the supernatant containing histones, was transferred to new Eppendorf tubes and dried using vacuum centrifugation. Histones were then resuspended with 0.2 M sulfuric acid. Following centrifugation, the histones in solution were transferred to new tube and subject to trichloroacetic acid precipitation for 5 minutes on ice. After centrifugation, the histones were washed with acetone and allowed to air-dry until moist. Histone pellets were resuspended in dd water, and protein concentration was determined using the Bradford protein assay.

### Histone Propionylation

10 µg of histone extract was resuspended in 20 μL of 50 mM ammonium bicarbonate and added to a 96-well plate for histone propionylation as adapted from Sidoli et al^46, 47^. The pH was monitored using Fisher pH test paper to be basic (pH 8.0). A 25% solution of propionic anhydride in acetonitrile was made and added to the histones at a 1:2 ratio of mixture:sample (v/v) to derivatize histone lysine and N-terminus. 10 μL of ammonium hydroxide was then added to the solution. After a 15-minute incubation, the samples were subject to a second round of propionylation by adding 20 µL of 25% propionic anhydride in acetonitrile and 20 µL of ammonium hydroxide. Using a TubroVap evaporation system, the plate was dried using vortex evaporation and heat. The histones were resuspended in 50 mM ammonium bicarbonate to a final concentration of 0.5 μg/μL. Trypsin (Promega) was added at a 1:50 trypsin:sample (w/w) ratio and allowed to incubate at room temperature overnight. The pH was checked to ensure a pH 8. Two rounds of derivatization were carried out to derivatize the tryptic peptide N-terminus.

### Peptide Desalting

Desalting was performed on derivatized peptides prior to LC-MS/MS analysis. First, stage tips were prepared following the protocol described in Lin et al^48^. Stage tips were activated with 50 µL of 100% methanol and centrifugation at 2000 g for 1 minute. Following activation, the stage tips were conditioned with 50 µL of 80% acetonitrile in water and centrifuged at 2000 g for 1 minute. Stage tips were then equilibrated with 50 μL of 0.1% trifluoroacetic (TFA) acid in water and centrifuged at 2000 g for 1 minute. Derivatized peptides for each condition were resuspended in 200 µL of 0.1% TFA in water and loaded onto the stage tips with centrifugation at 2000 g for 2 min. The stage tips were rinsed with 50 μL of 0.1% TFA in water with centrifugation at 2000 g for 2 min. Peptide trapped stage tips were transferred to a fresh 1.5 mL Eppendorf tube, and peptides were eluted twice with 75 μL of 80% acetonitrile in water with centrifugation at 2000 g for 1 min. Desalted peptides were vacuum centrifuged to dryness with no heat and stored at −20 °C until MS analysis.

### LCMS analysis

Derivatized and desalted peptides for each condition were resuspended in 0.1% FA in water for LCMS analysis. 200 ng of each sample was injected in an M5 MicroLC (SCIEX) coupled to a ZenoTOF 7600 mass spectrometer (SCIEX). An OptiFlow TurboV ion source was used with a microflow probe. Mobile phase A was 0.1% formic acid, and mobile phase B was 0.1% formic acid in acetonitrile. Samples were run by direct injection on a Kinetex XB C18, 100 Å, 2.6 μm, 0.3 x 150 mm column (Phenomenex) at a flow rate of 10 μL/min. The mass spectrometer was operated in ZenoSWATH DIA mode with CID fragmentation. The TOF MS scan range was m/z 120 to 1400. Dynamic collision energy was enabled and ranged from 14 to 46 V. TOF MS accumulation time was 100 ms, and MS2 accumulation time was 10 ms. The LC gradient and SWATH isolation scheme were optimized for a 10 min linear gradient from 5 to 32% solvent B.

### LCMS data analysis

Resulting Wiff files from each condition were converted to MZXML files using ProteoWizard MSConvert then searched on EpiProfile 2.1^49^ with EpiConvert extension against the human histone post translational modification (PTM) library with the label-free quantification filter applied. Microsoft Excel was used to conduct further data organization and Metaboanalyst 5.0^50^ was used to generate heatmaps for data visualization. No normalization, transformation or scaling was applied during data input. To visualize data in the heatmap, a Euclidean distance measure was applied with the Ward clustering method. All other parameters were default. Skyline^51^ was also used to perform manual validation of identified histone PTMs to ensure accuracy of data.

### IHC score

Automated scores were generated by image analysis. The slide was scanned and loaded into Qupath, representative areas of tumor, stroma, and necrosis were selectd, and a K-nearest neighbor classifier was created to assign cells to these categories. All cells were detected, classified as 0, 1+, 2+ or 3+ by DAB intensity, and classified them as tumor, stroma or necrosis. The h-score (column B) and Allred score (column E) were calculated only for tumor cells.

### Cell synchronization

A double-thymidine treatment (T/T; 18-h thymidine arrest and 8-h release followed by 18-h thymidine arrest) synchronized cells at the G1/S boundary. The thymidine concentration was 1mM.

### CHIP-seq data analysis

We mapped the levels of H3K9ac and H3K9me3 over *TP53, P63, P73, TOP1, TOP2A, TOP2B, ATM, KAP1*, and *NPAT* in several breast cancer cell lines using ChIP-seq data available on GEO (GSE85158)^52^. Briefly, the cell lines examined include the five major breast cancer subtypes including two ER-positive subtypes (Luminal-A and Luminal-B), the HER2-positive subtype, and two triple-negative subtypes (TNBC-Basal and TNBC-ClaudinLow). Additionally, normal-like immortal breast cells were included in the analysis. H3K9ac levels over these genes in the different breast cancer cell lines and normal-like cells follow a broadly similar pattern with some variations in their magnitude from GEO (GSE85158). The data was been normalized and sorted BAM files were converted to BigWig format using bamCoverage from the deepTools suite (v3.5.0). Normalization was applied using the Counts Per Million (CPM) method to account for differences in sequencing depth across samples. The conversion process included the use of a bin size of 10 bp and a smoothing window of 30 bp. The effective genome size was set to 2.7 billion bases to correspond to the mappable portion of the human genome. The Y axis is the same value for all the lines for easy comparison ≈0.48.

### Western blotting

Cells were lysed in RIPA buffer supplemented with protease and phosphatase inhibitors. Protein concentrations were determined using a BCA assay (Thermo Fisher). Equal amounts of protein (20–40 µg) were separated by SDS-PAGE and transferred to PVDF membranes (Millipore). Membranes were blocked in 5% non-fat dry milk in TBST (Tris-buffered saline with 0.1% Tween-20) for 1 hour at room temperature, followed by incubation with primary antibodies overnight at 4°C. After washing, membranes were incubated with HRP-conjugated secondary antibodies for 1 hour at room temperature. Protein bands were detected using enhanced chemiluminescence (ECL) and imaged with a Bio-Rad ChemiDoc system. Densitometry analysis was performed using ImageJ software.

## Data Availability Statement

The data supporting the findings of this study are available from the corresponding author upon reasonable request. All relevant datasets generated and/or analyzed during the current study will be made available in a publicly accessible repository upon publication.

## Abbreviations

H3K9AC: acetylation on histone H3 lysine 9
EGFR: Epidermal Growth Factor Receptor
HER2: Human Epidermal Growth Factor Receptor 2
ADCs: Antibody-drug conjugates
T-DXd: Trastuzumab deruxtecan
DSB: double-strand breaks
NCI ETCTN: National Cancer Institute Experimental Therapeutics Clinical Trials Network.

---

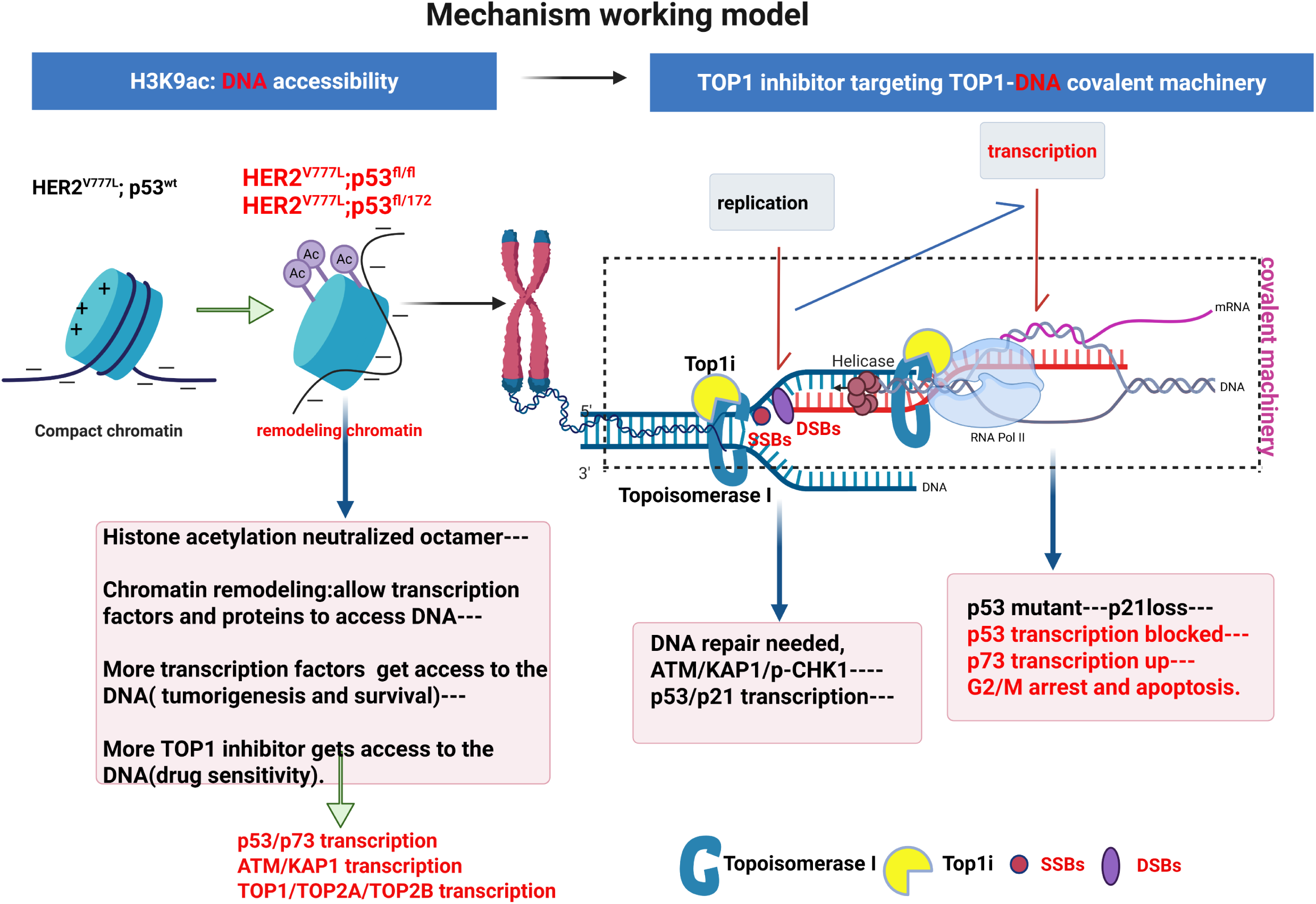

**Supplementary Fig. S1:**
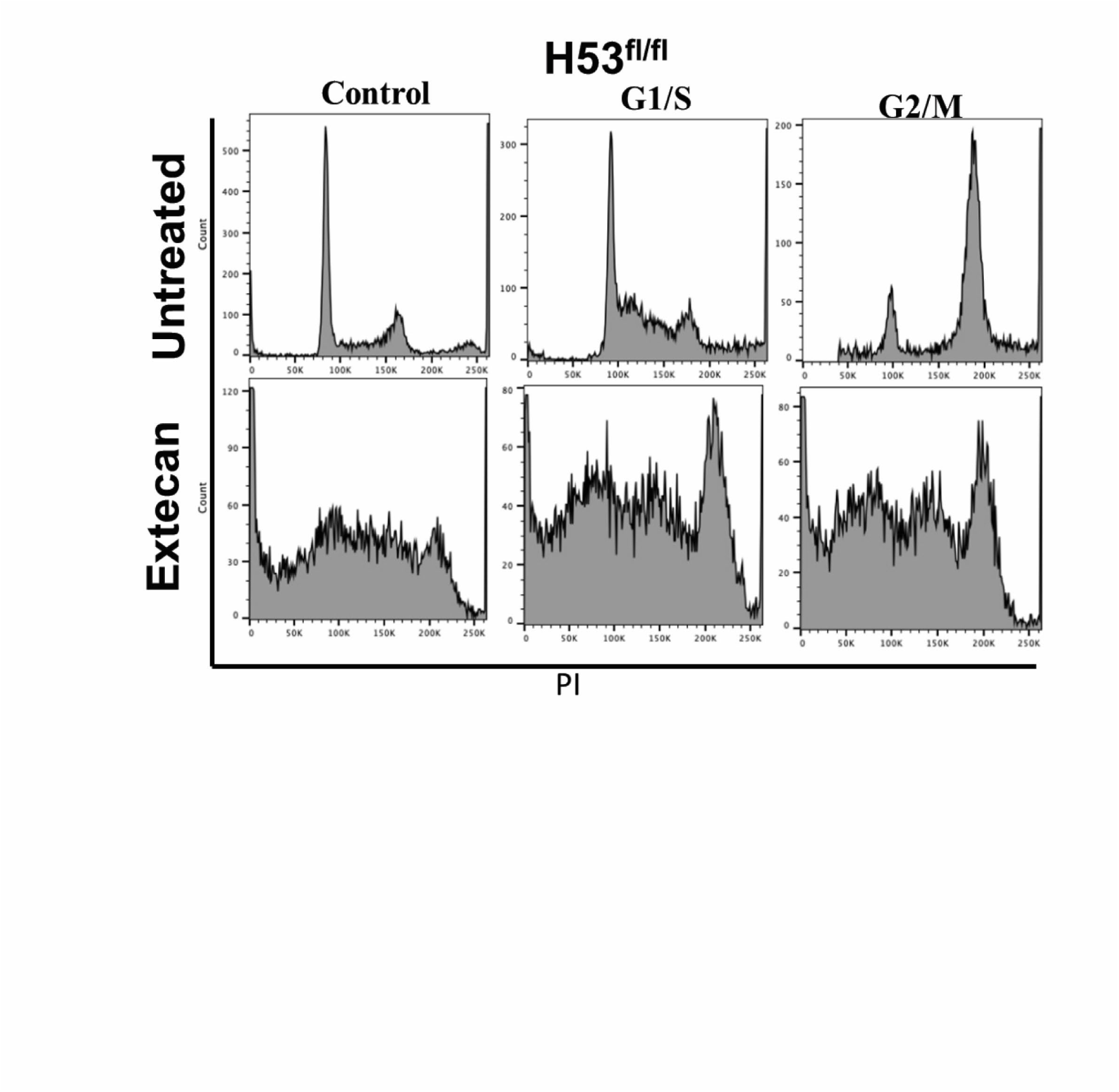
Cell cycle arrest causes mitotic catastrophe. Organoid cells are arrested at G1/S and G2/M phase by double thymidine treatment or thymidine-nocodazole treatment along with 100nM extecan treatment at the same time; PI staining shows the cell cycle status.

**Supplementary Fig. S2:**
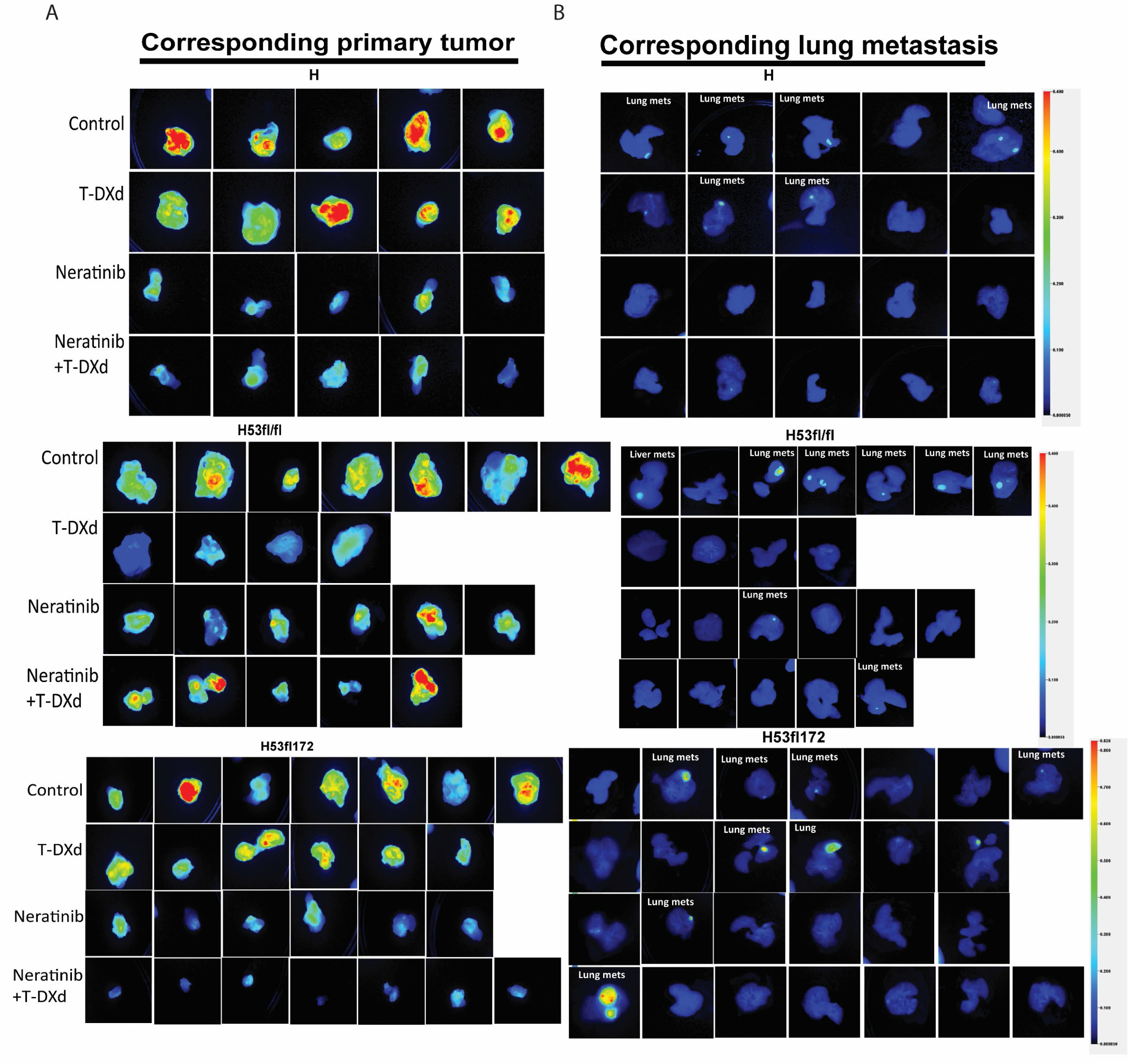
Lung metastasis in H53 mouse. (A) Imaging near-infrared fluorophore-labeled trastuzumab (NIR-trastuzumab) at 24 hours post-injection on the primary tumor of H, H53fl/fl and H53fl/172 mice in drug-treated groups as indicated. Neratinib was given daily at 40mg/kg by oral gavage. T-DXd was given weekly at 30mg/kg by tail vein injection for two weeks. (B) Imaging of near-infrared fluorophore-labeled trastuzumab (NIR-trastuzumab) at 24 hours post-injection on the metastatic lung tumors of H, H53fl/fl and H53fl/172 mice in drug-treated groups as indicated. Neratinib was given daily at 40mg/kg by oral gavage. T-DXd was given weekly at 30mg/kg by tail vein injection for two weeks.

**Supplementary Fig. S3:**
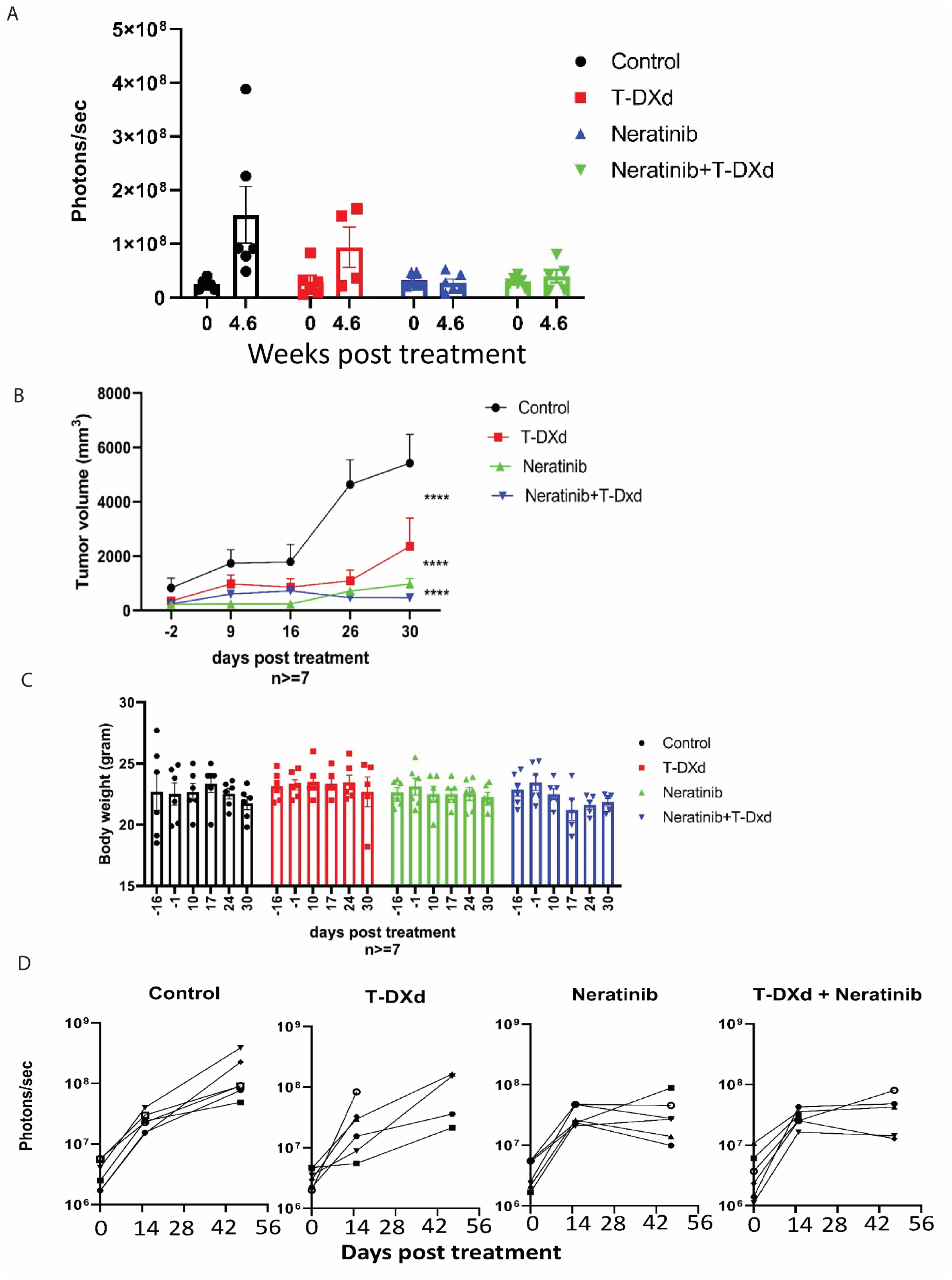
*In vivo* treatment results. (A) BLI intensity measurements of H53fl/fl organoid transplanted tumor growth luciferase signal on 4.6 weeks post-drug treatment. N>=6 per arm. (B) H53fl/fl organoid transplanted tumor volume was measured before or post-treatment as indicated. Data are plotted as means ± SEM. *P < .05, **P < .01, ***P < .001, and ****P < .0001 as calculated by the Unpaired t-test. (C) Mouse body weight was measured before or after treatment in H53fl/fl organoid transplanted mouse models. (D) BLI intensity measurements of H53fl/fl organoid transplanted tumor growth luciferase signal on each treated group indicated post-drug treatment.

**Supplementary Fig. S4:**
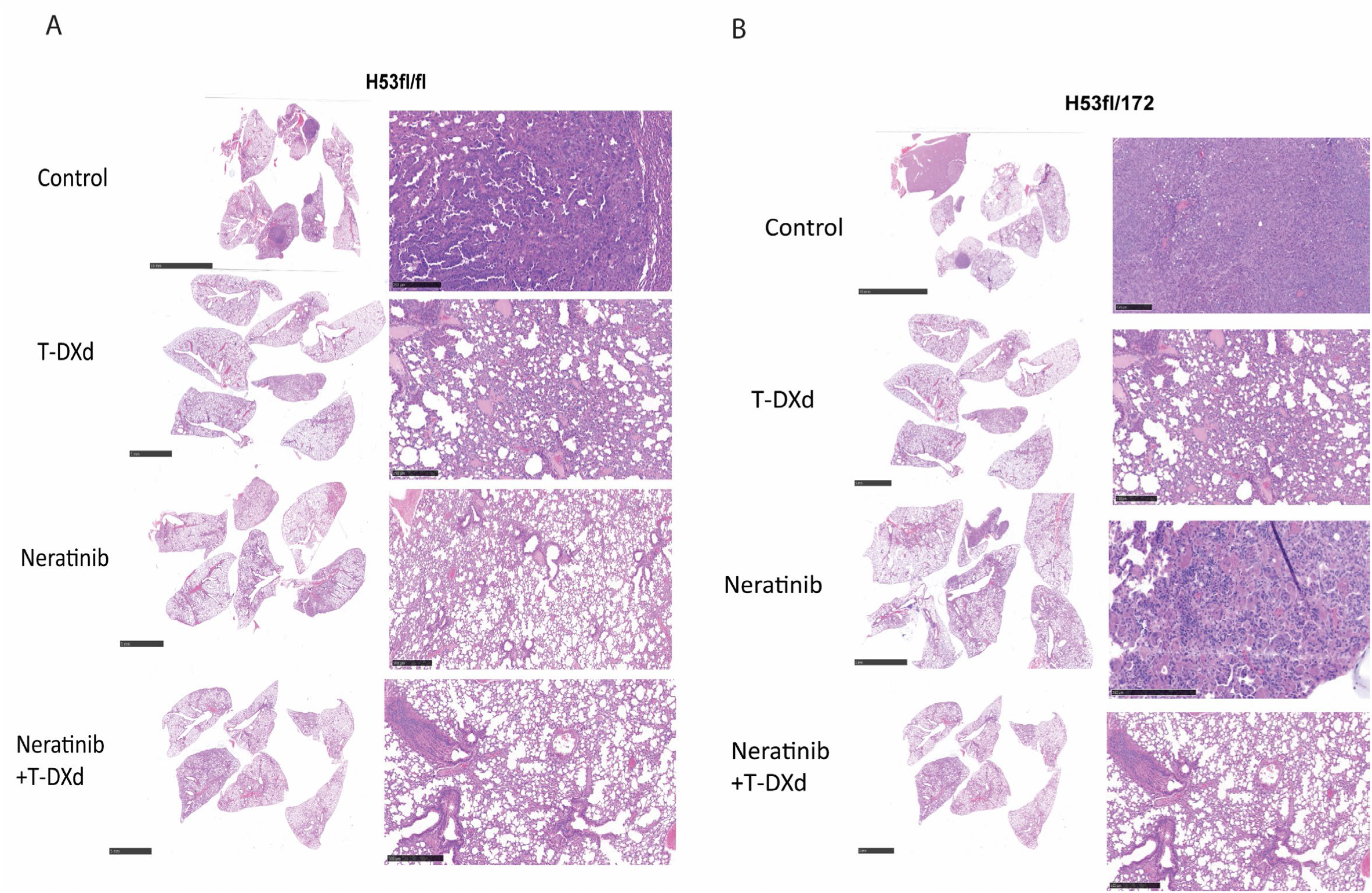
Lung metastasis morphology after treatment. (A) Representative hematoxylin and eosin (H&E) staining images of the lung from H53fl/fl transplant mice with or without treatment as indicated. Lower power scale bars are 5mm, high power scale bars are 500 μm. (B) Representative hematoxylin and eosin (H&E) staining images of the lung from H53fl/172 transplant mice with or without treatment as indicated. Lower power scale bars are 5mm, high power scale bars are 500 μm.

**Supplementary Fig. S5:**
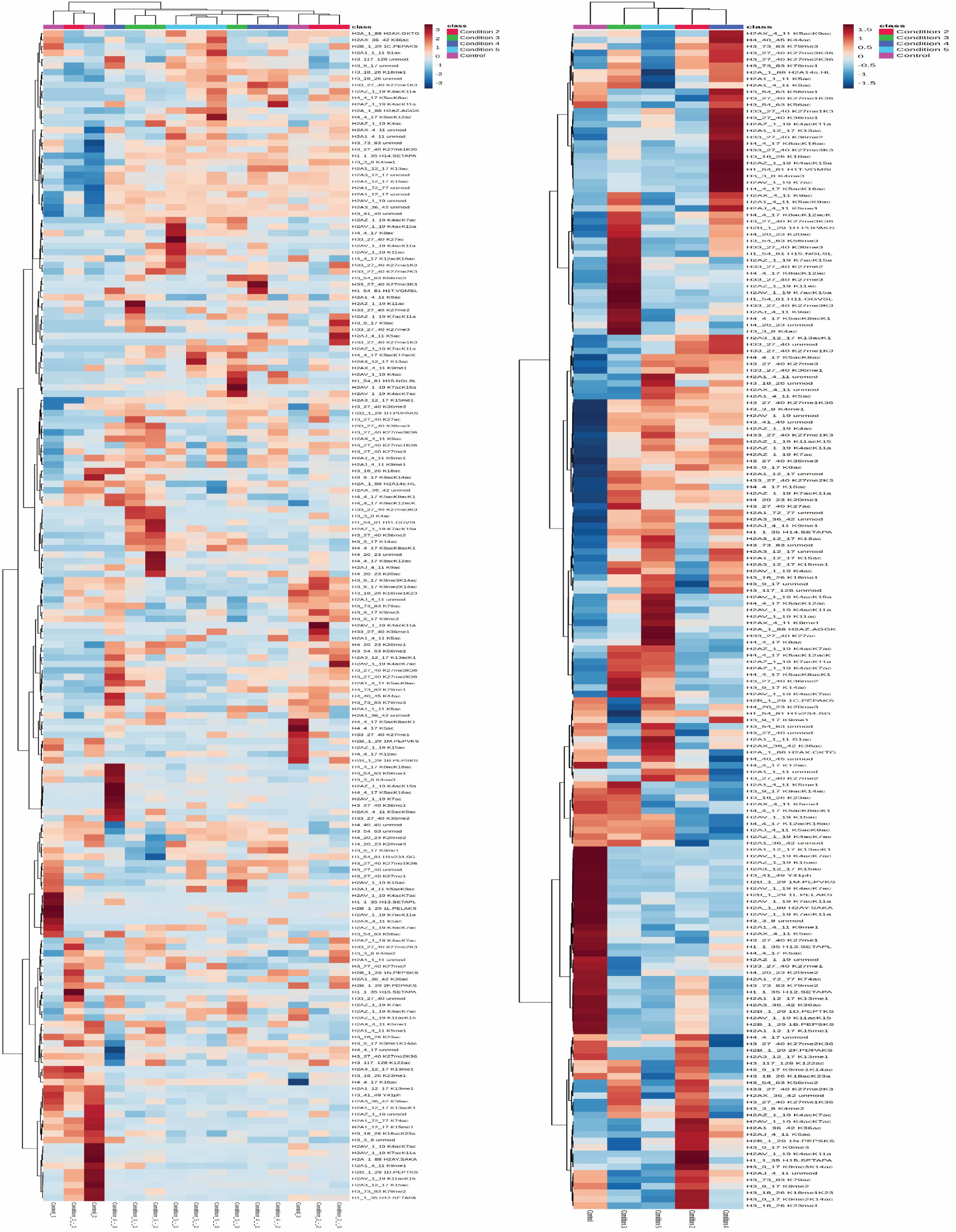
Histone modifications among each organoid cell group. Histone modifications heatmap with normalization and clusters by average ratios among each group of samples.

**Supplementary Fig. S6:**
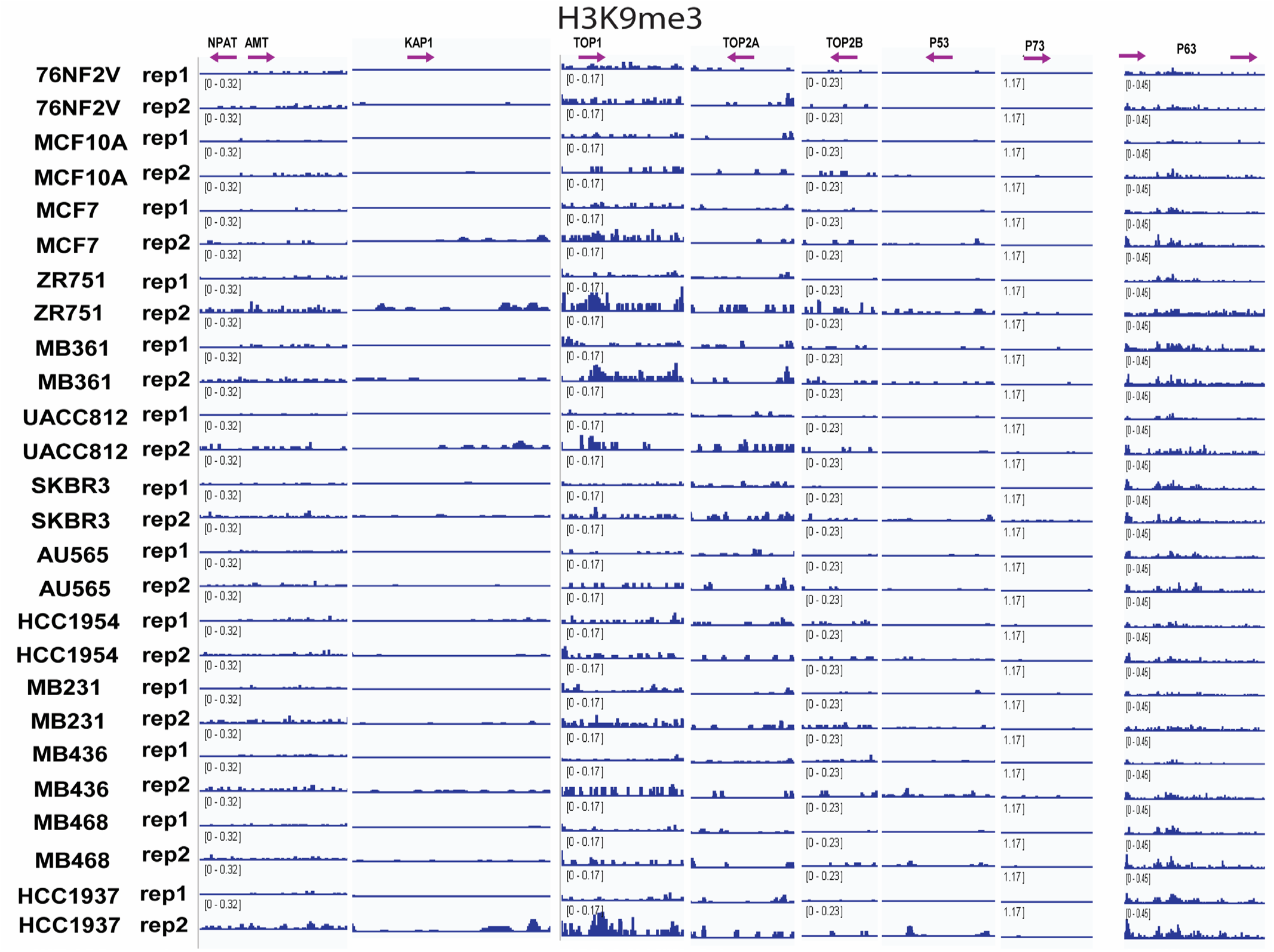
Location of H3K9me3 among the DNA repair-related factors in breast cancer.

## Notes

**Grant support:** This work was supported by the Department of Defense Breast Cancer Research Program grant number BC170330 (to R.B.), NIH/NCI PDXNet U54 CA224083 (to C.X.M. and R.B.), S10 OD027042 and S10 OD025264 (to the Mallinckrodt Institute of Radiology, Molecular Imaging Center), and the Siteman Cancer Center (SCC) Support Grant P30CA091842.

**Conflict of Interest Disclosures:** R. Bose received a research grant from Puma Biotechnology, Inc. and has performed consulting on a HER2 clinical trial for Genentech. Neratinib used in this study was provided by Puma Biotechnology, Inc.

### Competing Interest Statement

R. Bose received a research grant from Puma Biotechnology, Inc. and has performed consulting on a HER2 clinical trial for Genentech. Neratinib used in this study was provided by Puma Biotechnology, Inc.

